# Bacteriophage-host interactions in microgravity aboard the International Space Station

**DOI:** 10.1101/2023.10.10.561409

**Authors:** Phil Huss, Chutikarn Chitboonthavisuk, Anthony Meger, Kyle Nishikawa, R.P. Oates, Heath Mills, Olivia Holzhaus, Srivatsan Raman

## Abstract

Bacteriophage-host interactions play a fundamental role in shaping microbial ecosystems. While extensively studied on Earth, their behavior in microgravity remains largely unexplored. Here, we report the dynamics between T7 bacteriophage and *E. coli* in microgravity aboard the International Space Station (ISS). Phage activity was initially delayed in microgravity but ultimately successful. We identified *de novo* mutations in both phage and bacteria that enhanced fitness in microgravity. Deep mutational scanning of the phage receptor binding domain revealed striking differences in the number, position, and mutational preferences between terrestrial and microgravity conditions, reflecting underlying differences in bacterial adaptation. Combinatorial libraries informed by microgravity selections yielded T7 variants capable of productively infecting uropathogenic *E. coli* resistant to wild-type T7 under terrestrial conditions. These findings help lay the foundation for future research on the impact of microgravity on phage-host interactions and microbial communities and the terrestrial benefits of this research.

## Introduction

The interaction between bacteriophages (or ‘phages’) and their bacterial hosts plays a fundamental role in shaping microbial ecosystems both in humans and in the environment. These interactions are influenced by the physical forces of fluid mixing and the underlying physiology of the bacterial host and the phage. Although phage-host interactions have been extensively studied in terrestrial ecosystems, the impact of microgravity on these interactions has yet to be fully investigated. Studying phage-host interplay in microgravity may reveal new mechanisms with relevance both in space and on Earth.

Microgravity presents physical and physiological challenges to both phage predation and bacterial growth. Physically, gravity influences phage movement and their ability to encounter bacterial hosts. Phages diffuse randomly in fluid until they contact a bacterial cell, after which van der Waals forces and electrostatics drive adsorption [1,2]. On Earth, gravity induces natural convection, enhancing fluid mixing. Buoyancy and sedimentation caused by gravity move phages and nutrients toward bacterial cells. In microgravity, materials of differing densities disperse evenly, disrupting nutrient diffusion and bacterial motility [3–6]. Convection, driven by temperature-dependent changes in density, requires gravity to occur [5,7]. The absence of gravity profoundly impacts bacterial metabolism and stresses bacteria [7–9]. Microgravity increases biofilm formation and metabolic rates [10,11], while reduced mixing limits waste removal and access to nutrients. This induces the overexpression of starvation genes and increases membrane flux. Bacteria may adapt by altering their proteome, including host factors such as receptors that are essential for phage infection. In summary, microgravity is a distinct environmental niche that may significantly alter phage-host dynamics, with broad implications for microbial communities in space.

In this study, we investigated how microgravity affects interactions between T7 bacteriophage and non-motile *E. coli* BL21 aboard the International Space Station (ISS). Our evaluation of short-term (hours) and long-term (23 days) incubation of phage and host in microgravity showed significant differences in phage and bacterial viabilities and phage activity compared to terrestrial controls. Phages accumulated many *de novo* mutations over time that may enhance receptor binding or phage infectivity, while bacteria acquired *de novo* mutations in genes that may enhance fitness in microgravity and counter phage predation. Deep mutational scanning (DMS) of the phage receptor binding protein (RBP) in microgravity revealed a fitness landscape significantly different from our terrestrial experiments, suggesting substantial differences in the host receptor profile and selection pressure under microgravity. Notably, a combinatorial library of RBP variants enriched in microgravity exhibited a significant improvement in activity against terrestrial uropathogenic E. coli, while a similar library derived under terrestrial conditions showed no gain, highlighting microgravity as a source of insights into phage-host dynamics with relevance to Earth. Overall, our findings help lay a foundation for future research into the impact of phage-host interactions on microbial communities in microgravity and in the context of space exploration.

## Results

### Design of experiments for the International Space Station

We prepared two identical sets of 32 sealed cryovial tubes containing experimental samples: one designated for incubation in microgravity, and the other designated for terrestrial incubation (Figure 1). Each set was divided into four prepackaged groups of eight tubes for incubation at 37°C. Three groups were incubated for short-term time points (1, 2, and 4 hours), and one group for a long-term time point (23 days).

**Figure 1.**
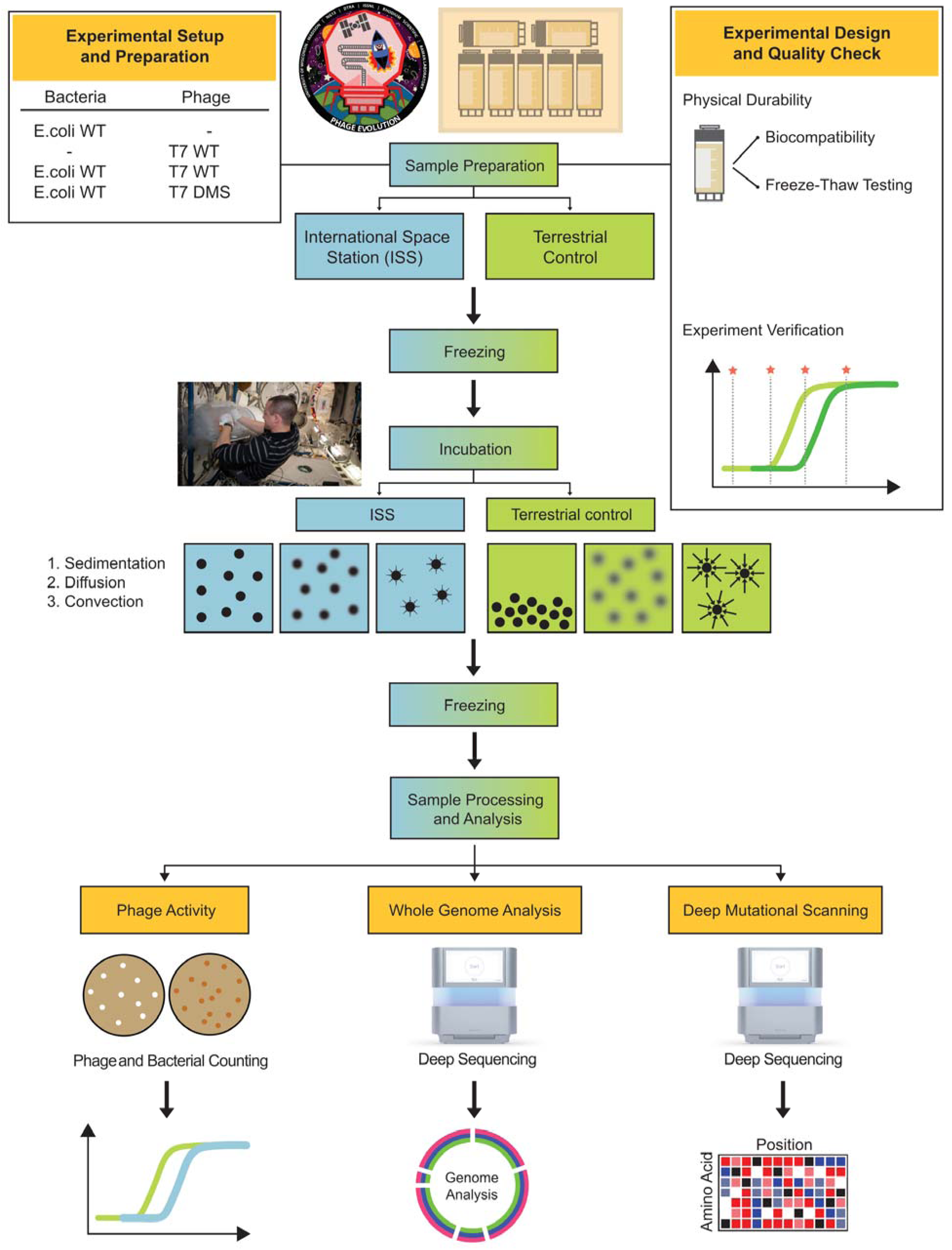
Experimental design to evaluate microgravity interactions on the ISS. Samples were prepared on Earth, with quality checks to ensure cryovial integrity and prevent leakage during freeze–thaw cycles. Identical sets were frozen, then thawed and incubated either in microgravity on the ISS (left) or terrestrially (right) for defined intervals. All samples were re-frozen and later analyzed on Earth for phage and bacterial titers, whole-genome sequencing, and deep mutational scanning of the T7 receptor binding protein tip domain. *Image courtesy of NASA/Rhodium Scientific*.

Each short-term group included three replicates of T7 and *E. coli* BL21 mixed at a phage to host ratio (multiplicities of infection or MOIs) of 10^-6^ and 10^-4^ and two sample controls with T7 phage only or *E. coli* only. All bacterial samples contained 4 mL of log-phase (OD_600_ ∼0.4) *E. coli* with an estimated titer of ∼1-2×10^8^ CFU/mL. The initial bacterial concentration and MOI were selected to allow for measurable changes in phage titer after incubation, accounting for anticipated cell and phage viability loss. To isolate the effects of microgravity, we used a non-motile *E. coli* strain, removing the variable of host motility enhancing fluid mixing. The 23-day group included three replicates of T7 phage and *E. coli* mixed at an MOI of 10^-4^, three replicates of T7 DMS library mixed with *E. coli* at an MOI of 10^-2^, and two control samples of T7 only and *E. coli* only. The higher MOI for the DMS group compensated for the lower abundance of individual variants. The DMS library comprises 1,660 T7 variants, each with a single amino acid substitution in the tip domain of the receptor binding protein (RBP), that has been previously tested under terrestrial conditions [12].

The cryovial containers passed biocompatibility, leak testing, and experimental validation (see Supplementary File 1) to ensure sample integrity and comply with NASA safety standards. All samples were prepared on Earth by mixing phage and bacteria in Rhodium cryovials and immediately freezing them at −80°C. Frozen samples were shipped to NASA’s Wallops Flight Facility 24 days before launch and transported to the ISS aboard the Northrop Grumman NG-13 Cygnus rocket. After incubation in microgravity, the samples were refrozen, transported back to Earth, and delivered to our laboratory. We then thawed the samples, measured phage and bacterial titers, sequenced their genomes, and analyzed the DMS library (Figure 1). We recorded the duration of freezing and incubation aboard the ISS and evaluated the second set of samples terrestrially using the same incubation and freezing times.

### Bacteriophage T7 activity is reduced in microgravity

Under normal terrestrial conditions with shaking at 37°C, the T7 phage infects and lyses *E. coli* BL21 within 20-30 minutes and produces 100-200 progeny phages [13,14]. We hypothesized that in microgravity, reduced fluid mixing would slow the infection cycle by limiting productive encounters between phages and bacteria. Additionally, microgravity-induced stress might disrupt host homeostasis, alter receptor expression, or interfere with intracellular processes, impeding successful phage replication. To test this hypothesis, we measured phage and bacterial titers after 1-, 2-, and 4- hours, as well as after 23 days of incubation. Because the extent of phage replication delay in microgravity was unknown, this approach allowed us to capture a broad range of possible delays.

Phage and bacteria were co-cultured at MOIs of 10^-6^ and 10^-4^ for short-term incubation time points (1-, 2- and 4-hours) and 10^-4^ for long-term incubation (23-days). A significant increase in phage titer and decrease in bacterial host titer would indicate successful phage activity. Under terrestrial conditions phage titers increased significantly by 5-7 logs and bacterial titers decreased significantly by 4-5 logs after four hours, regardless of the MOI (Figure 2A). Phage infection thus occurred between two and four hours for the terrestrial samples, indicating the experimental conditions delayed the infection cycle by approximately two hours while allowing for successful phage replication.

**Figure 2.**
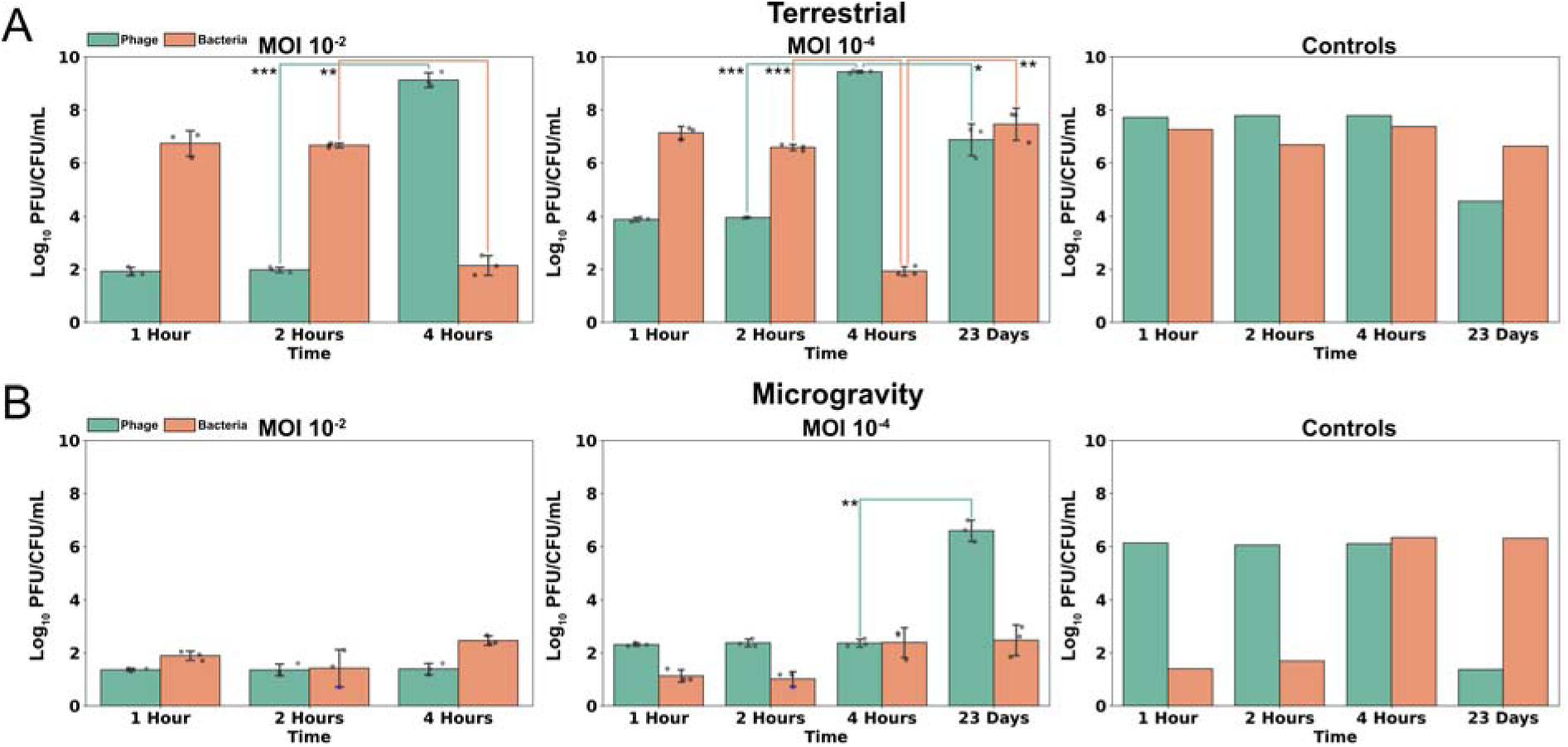
Bacteriophage T7 growth is inhibited by microgravity. The titer of phage (green) and bacteria (orange) samples (log_10_ Plaque Forming Units or Colony Forming Units, PFU/CFU/mL) after **(A)** terrestrial incubation or **(B)** incubation in microgravity mixed at an MOI of 10^-2^ (left), 10^-4^ (middle), or incubated separately as controls (right). Bars show mean ± SD; triplicate samples shown as points, with blue indicating values at the limit of detection. The significance between adjacent time points was assessed by two-sample t-test (*p<0.05, **p<0.01, ***p<0.001).

Under microgravity conditions, we observed no increase in phage titer at any short-term incubation time points at either MOI, but a significant 4-log increase at the 23-day time point (Figure 2B). This result indicates microgravity did not prevent productive infection and lysis but delayed it to some point past the four-hour timepoint. The persistence of bacteria at the 23-day time points (approximately 10^7^ CFU/mL for the terrestrial samples and 10^2^ CFU/mL for microgravity samples) also suggests that a phage-resistant bacterial population emerged in both conditions.

Bacterial titer without phage remained stable at early time points under terrestrial incubation but fell 6-7 logs at the 1- and 2-hour time points in microgravity. We postulate that microgravity-induced stress impaired the bacteria’s ability to survive subsequent freezing and bacteria were able to recover by the 4-hour time point. Phage without bacteria saw a 2-log decrease in viable titer in early time points in microgravity compared to terrestrial incubation but otherwise appeared more stable than bacteria, except for the 23-day timepoint, where we observed a 4- and 7-log fold decrease in phage titers terrestrially and in microgravity respectively. Phages are known to lose viability over time without a propagating host [15,16], and this effect appeared more pronounced in microgravity.

These experiments demonstrate that microgravity challenges both phage and bacterial viability. While phage infectivity is delayed compared to terrestrial conditions, phages ultimately overcome this barrier and successfully infect their hosts. Future studies targeting intermediate time points will be critical for defining the precise latent period under microgravity.

### Enriched mutations are distributed broadly in T7 phage

Next, we sought to identify mutations in the phage or bacterial genome that influenced phage-host interactions under microgravity. We performed whole-genome sequencing of T7 and *E. coli* BL21 before and after incubation, using pre-incubation genomes as references to identify *de novo* mutations in the 23-day samples from each condition. To determine whether *de novo* non-synonymous substitutions or frameshifts in T7 were significantly enriched, we compared the pooled frequencies of abundant non-synonymous mutations to the distribution of synonymous *de novo* substitutions in each condition (Mann-Whitney U test, FDR-adjusted p<0.05; Figure 3A, Supplementary File 2). To assess whether specific genes had significantly more non-synonymous substitutions than other genes, we calculated this mutation density for each gene and compared it to the average mutation density per condition (one-tailed t-test, FDR-adjusted p<0.05, one-sided 95% CI; Figure 3B, Supplementary Figure 1). Finally, we compared the gene-level distribution of non-synonymous mutations between microgravity and terrestrial conditions (Mann-Whitney U test, FDR-adjusted p<0.05) to identify genes with condition-specific enrichment of these mutations (Figure 3C, Supplementary Figure 2).

**Figure 3.**
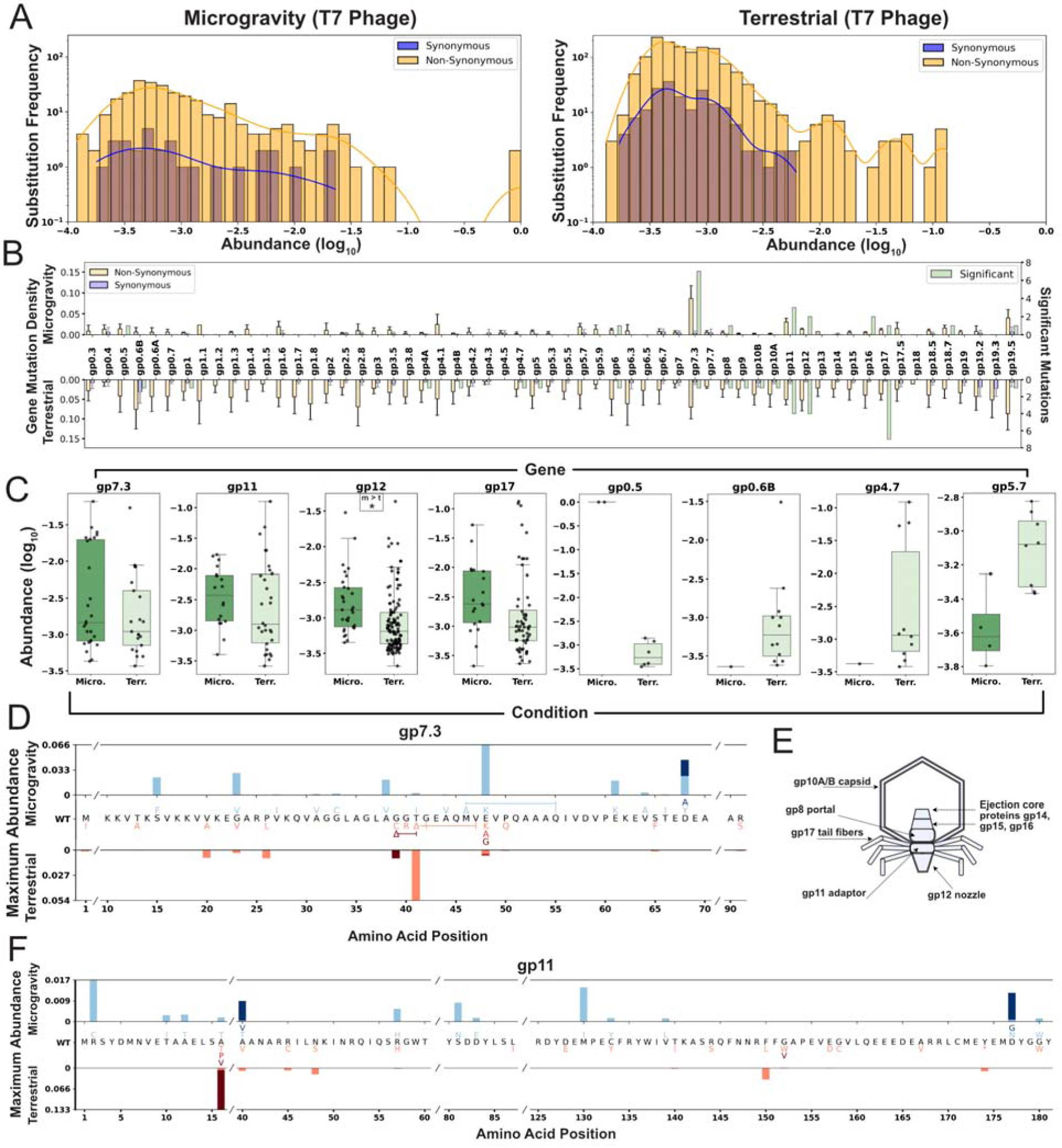
Enriched mutations are distributed broadly in T7 phage. **(A)** Substitution frequency and abundance of phage *de novo* synonymous (blue) and non-synonymous substitutions or frameshifts (yellow) after microgravity (left) or terrestrial (right) incubation. **(B)** Average gene mutation density ±SD for *de novo* synonymous (blue) and non-synonymous (yellow) substitutions after microgravity (top) and terrestrial (bottom) incubation. Green bars show the number of significant mutations per gene from (A). **(C)** Log_10_ abundance of mutations in selected phage genes after microgravity (left, dark green, Micro.) and terrestrial (right, light green, Terr.) incubation. Significance shown as * (adjusted p<0.05, one-tailed t-test with FDR) **(D)** Maximum substitution abundance in *gp7.3* after microgravity (top, blue shading) or terrestrial (bottom, red shading) incubation. Wild-type (WT) sequence shown center with substitutions shaded matching bar color. Deletions shown as Δ; slashes (/) in WT sequence mark omitted unmutated regions. **(E)** Diagram of selected structural genes. *Gp7.3* is excluded due to an uncertain structural role. **(F)** Maximum substitution abundance in *gp11*, displayed as in (D).

Significantly enriched (p<0.05) phage substitutions were found across both structural and non-structural proteins under terrestrial and microgravity conditions (Figure 3B). In microgravity, gene product (*gp*) 7.3 and *gp11* exhibited significantly more *de novo* non-synonymous substitutions than other genes (Figure 3B, Supplementary Figure 1A). Mutation density was overall higher terrestrially and no gene showed significant enrichment compared to others terrestrially (Supplementary Figure 1B). Although *gp7.3* is not fully characterized, it is considered essential for T7 infectivity in *E. coli* BL21 under terrestrial conditions [17]. This small 99-amino-acid protein may function as a scaffolding protein or contribute to host adsorption, though its role in the mature virion remains uncertain [18–20]. *gp7.3* harbored seven significantly enriched substitutions in microgravity, the highest number observed in any gene under that condition. These substitutions were distributed throughout the protein (Figure 3D, Supplementary Figure 3), with four notable changes (E48K, E61K, D68Y, D68A) involving substantial shifts away from negatively charged residues. The only significantly enriched mutation in *gp7.3* terrestrially was a six-amino-acid deletion spanning G42 to V47. The region from G39 to Q50 contained a dense cluster of substitutions and in-frame deletions, including a 3-amino-acid deletion (G39-T41) terrestrially and a deletion from M46 to Q55 in microgravity, all occurring in a region of the protein predicted to be unstructured (Supplementary Figure 3). The high number of enriched substitutions and recurring in-frame deletions in this small protein suggest that *gp7.3* is both structurally flexible and critical for phage activity in both environments.

*gp11* is an adaptor protein within the T7 tail that connects the portal protein *gp8*, the nozzle protein *gp12*, and the six subunits of the tail fiber protein *gp17* (Figure 3E) [17,19,21]. Enriched substitutions were distributed throughout *gp11*, spanning both exposed and buried residues (Figure 3F, Supplementary Figure 4A–B). One significantly enriched substitution, R2C, arose independently twice in microgravity and is located in a flexible region capable of directly interacting with *gp17* tail fibers (Supplementary Figure 4C–D). These findings suggest that the substitutions may influence phage fitness by altering *gp11*’s structure or stability rather than through direct interaction with the bacterial host.

Comparison of mutation abundance revealed that *de novo* non-synonymous substitutions were significantly more prevalent in the nozzle protein *gp12* after incubation in microgravity than under terrestrial conditions, suggesting a more prominent role for this protein in microgravity (Figure 3C). Of the six individually enriched non-synonymous substitutions identified across both conditions, five involved changes toward positively charged residues (Q184R, R205H, Q242R, K404R, and W707R) (Supplementary File 2). These substitutions were distributed throughout the protein, with three more likely contributing to host interactions (Supplementary Figure 5). Specifically, R205 is surface-exposed and positioned near the host, Q242 lies close to the terminus of the DNA delivery channel, and Q184 faces directly toward the host. The charge shifts and spatial distribution of these substitutions highlight the functional importance of *gp12* in enhancing phage fitness under both terrestrial and microgravity conditions.

Several other significantly enriched substitutions were particularly notable. In microgravity, the V26I substitution in *gp0.5* was the only mutation to sweep the entire phage population—and did so independently in two replicates—indicating a strong fitness advantage. *gp0.5* is an uncharacterized class I gene, potentially associated with the host membrane due to the presence of a putative transmembrane helix [22]. Under terrestrial conditions, the T115A substitution in *gp4.7* was significantly enriched and highly abundant across all three replicates. No mutations were detected in this gene under microgravity, suggesting selection pressure may be unique to terrestrial conditions. Although the function of gp4.7 remains unknown, BLASTP analysis identified homologs with ∼40% similarity to putative HNH endonucleases in *Klebsiella* and *Pectobacterium* phages [22].

Lastly, numerous significantly enriched substitutions were found in the tail fiber *gp17*, particularly under terrestrial conditions (Figure 3B). In both environments substitutions were concentrated in the C-terminal tip domain, with repeated mutations at D540 and neighboring residues. This region is a known determinant of host range and infectivity in terrestrial *E. coli* strains [12], and these results suggests continued importance during prolonged incubation in both gravity conditions.

### Enriched bacterial mutations reflect phage-mediated selection

*De novo* mutations in *E. coli* BL21 were significantly more abundant in samples mixed with phages than in those without phages under both terrestrial and microgravity conditions, indicating strong phage-driven selective pressure in both environments (Figure 4A, Supplementary Figure 6A-C, Mann-Whitney U test p<0.001, Kaplan-Meier survival and log-rank statistical test p<0.001). Significantly enriched non-synonymous *de novo* bacterial substitutions and frameshifts were present in both conditions (Figure 4B, Mann-Whitney U test, FDR-adjusted p<0.05, >25% abundance, Supplementary File 3), and pooling genes on Gene Ontology (GO) categories [23,24] revealed that most enriched genes were associated with membrane function or the regulation of metabolic process (Figure 4C).

**Figure 4.**
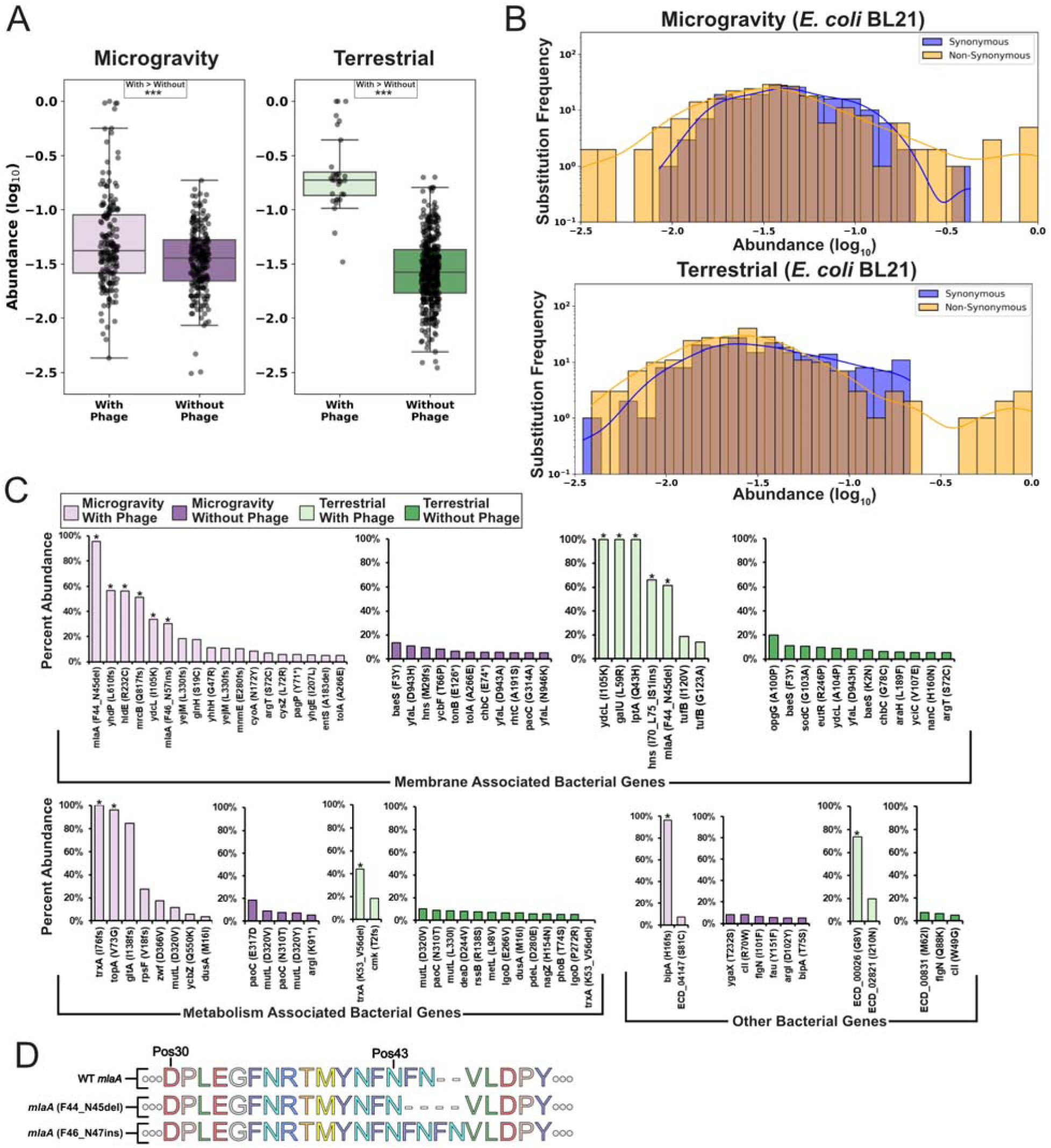
Enriched bacterial mutations reflect phage-mediated selection. **(A)** Boxplots of log_10_ abundance for *de novo* non-synonymous substitutions and frameshifts after microgravity (left) or terrestrial (right) incubation, comparing incubation with (light shading) and without (dark shading) phage. Significance assessed by Mann Whitney U test (***p<0.001), with > indicating the more abundant population. **(B)** Frequency for bacterial *de novo* synonymous (blue) and non-synonymous substitutions or frameshifts (yellow) after microgravity (top) or terrestrial (bottom) incubation. **(C)** Maximum abundance of *E. coli* BL21 non-synonymous substitutions or frameshifts (>5%) after incubation in microgravity with phage (light purple) or without phage (dark purple), and after terrestrial incubation with phage (light green) or without phage (dark green). Grouped by membrane-associated genes (top), metabolism-associated genes (bottom-left), or other genes (bottom-right). Mutations significantly enriched in (B) marked with stars. **(D)** Illustration of *mlaA* mutation effects. Deletions and insertions result in loss or repetition of phenylalanine and asparagine residues.

Bacterial mutations significantly enriched (p<005) only under microgravity were frequently associated with the outer membrane and cellular stress response. Notable examples include *hldE* (56.5%, R232C) associated with the synthesis of the lipopolysaccharide (LPS) core [25]; *mrcB* (51.6%, Q817 frameshift), which plays a role in cell wall synthesis and permeability [26,27]; and *bipA* (96.5%, H16 frameshift, also known as *typA*) linked to LPS biosynthesis and temperature sensitivity, and previously associated with truncated LPS phenotypes [35–37]. The significant abundance of this mutation suggests that *bipA* might play a role in phage sensitivity in microgravity. *topA*, a DNA topoisomerase associated with stress response, was also significantly enriched in microgravity (96.1%, V73G) [28,29]. Additional mutations unique to microgravity included *gltA* (85%, I138 frameshift), a citrate synthase [30] and *rpsF* (27.7%, V18 frameshift), which encodes a 30S ribosomal protein [31]. Under terrestrial conditions, bacterial mutations significantly enriched in the presence of phage included *galU* (100%, L59R), involved in UDP glucose metabolism and associated with O-polysaccharide in other strains [32,33]; *lptA* (100%, Q43H), responsible for LPS assembly [34]; and *hns* (66%, I70_L75 IS1 insertion), a global DNA-binding protein responsible for regulating metabolism and nutrient acquisition [35].

Several genes had significantly enriched mutations under both terrestrial and microgravity conditions. *trxA* is a processivity factor for T7 DNA polymerase and is a known essential gene for phage activity [36,37]. Deletions in *trxA* were significantly enriched in both conditions (terrestrial: 44.4%, K53_V56d deletion, microgravity: 100%, I76 frameshift), indicating the gene remains essential to the phage in microgravity. The same substitution in *ydcL* was significantly enriched in both conditions (I105K, terrestrial 100%, microgravity 33.9%). *ydcL* encodes a transcriptional regulator that triggers small, slow-growing persistor cell states, which could benefit bacteria during prolonged incubation conditions like those in this experiment [38,39].

Finally, intriguing indels were significantly enriched in *mlaA* in both conditions (Figure 4D). In each condition, two samples exhibited 6-bp deletions resulting in the loss of amino acids F44 and N45 (microgravity abundance: 95.5% and 16.5%; terrestrial abundance: 61.5% and 24.9%). In contrast, the third microgravity sample showed significant enrichment of a 6-bp insertion that added Asp and Phe, after N45 (30.2%, F46_N47ins), effectively inserting and repeating the same two amino acids deleted in the other samples.

mlaA encodes an outer membrane lipoprotein believed to remove mislocalized phospholipids from the outer membrane and shuttle them back to the inner membrane [40]. This gene has not yet been associated with changes in phage activity. A mutant with the same F44_N45 deletion has been characterized in *E. coli* MC4100 [41,42]. This mutation increases outer membrane permeability, phospholipid accumulation, and vesiculation—changes that could affect phage adsorption and potentially confer a competitive advantage. A prior study found that this deletion eventually led to bacterial cell death [42], but our results suggest this mutation may enhance bacterial survival under phage pressure. This discrepancy could also reflect differences in selection context, strain background, or the presence of suppressor mutations. Supporting this possibility, we also identified a significantly enriched frameshift-inducing deletion in *yhdP* (56.9%, L610 frameshift) in a microgravity sample that had the most abundant *mlaA* deletion. *yhdP* is involved in phospholipid transport to the outer membrane, and its loss has been shown to slow transport and reduce cell death in F44_N45 *mlaA* mutants [41], suggesting it may similarly enhance survivability in microgravity.

### Deep Mutational Scanning profiles beneficial substitutions in microgravity

Bacteria often resist phage predation by mutating or downregulating surface receptors essential for phage adsorption. Microgravity-induced stress may amplify this response, altering the bacterial proteome—including phage receptor profiles. Such changes can drive adaptive mutations in the phage receptor binding protein (RBP). To investigate these interactions, we examined how individual substitutions in the tip domain of the T7 RBP affect phage viability in microgravity.

The T7 RBP consists of six short non-contractile tails that form a homotrimer composed of a rigid shaft ending with a β-sandwich tip domain [43]. This domain is a key determinant of host recognition and interacts with host receptor LPS to position the phage for successful, irreversible binding [37,44–48]. We conducted comprehensive single-site saturation mutagenesis of the RBP tip domain, generating a library of 1,660 variants spanning residues 472–554 (based on PDB 4A0T). We then sequenced and compared mutational enrichment profiles following the 23-day selection under terrestrial and microgravity conditions.

We recovered phage DNA from each sample and scored each variant based on its relative abundance before and after selection (functional score, *F*) normalized to wildtype (normalized functional score, *F_N_*). Scores were averaged across replicates, and only variants present in at least two replicates were retained for analysis. Although significant dropout of low-performing variants was expected due to the extended incubation, we successfully determined scores for 51.2% (880) of variants in microgravity and 39% (648) in terrestrial conditions (Figure 5A–B, Supplementary Figure 7A, Supplementary File 4). Variant scores correlated well across replicates despite differences in phage titer and reflected multiple rounds of replication over the 23-day incubation period, suggesting that lower-titer samples underwent selection but subsequently lost viability (Supplementary Figure 7B-D). On average variants had significantly higher scores after terrestrial incubation (two-sample t-test, Mann-Whitney U, p<0.001) (Supplementary Figure 8A) with the wild-type phage performing significantly worse terrestrially compared to microgravity (terrestrial *F* = 0.58, microgravity *F* = 3.5, p<0.01).

**Figure 5.**
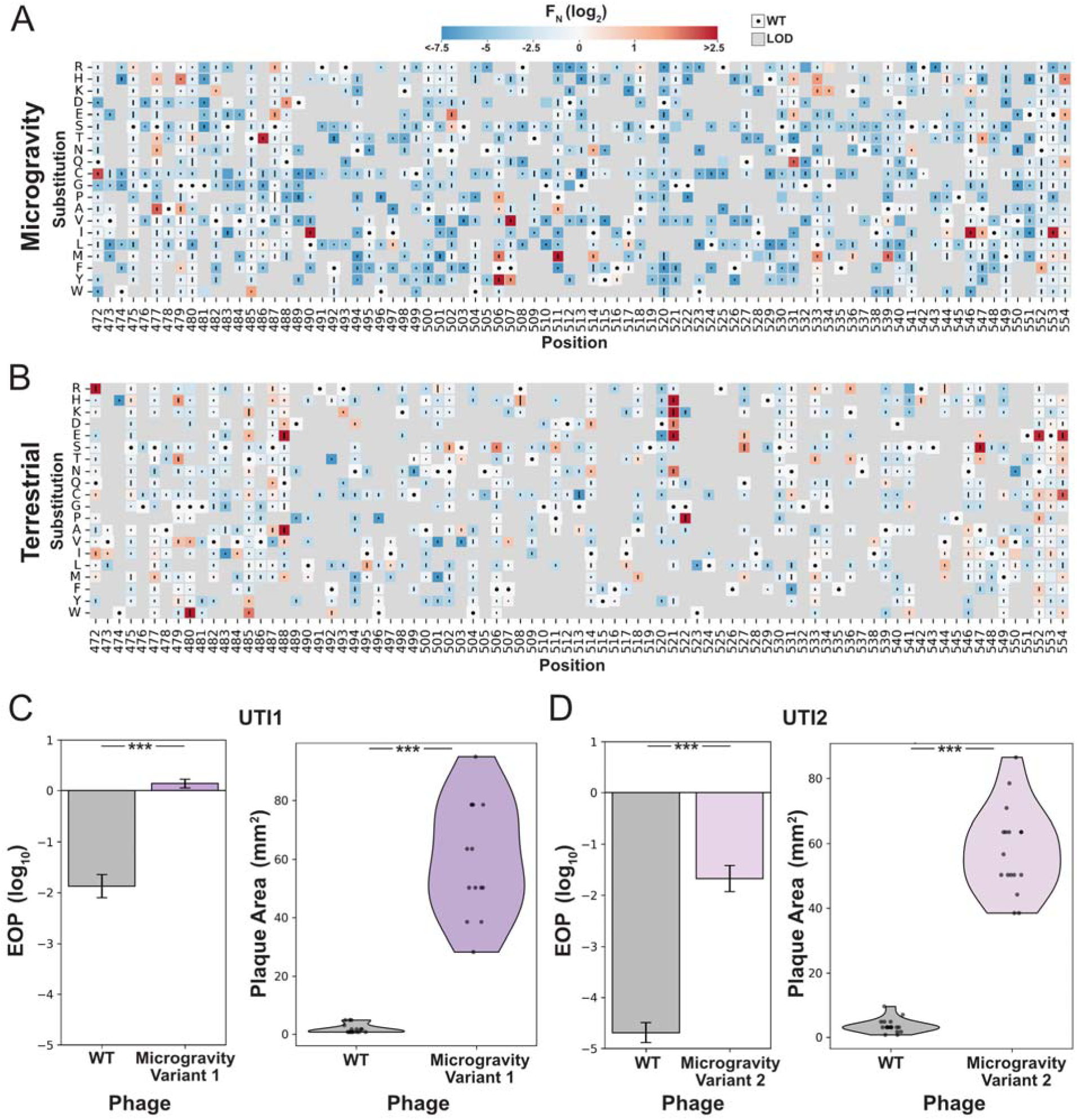
Deep Mutational Scanning reveals beneficial substitutions in microgravity. **(A-B)** Heatmaps of normalized functional scores (F_N_ log_2_) for all RBP substitutions after microgravity (A) and terrestrial (B) incubation. Scores shown on a blue to red gradient; Wildtype (WT, *F_N_* log_2_ =0) in white with a black dot; variants below the limit of detection (LOD) in grey. Line length represents standard deviation. Substitutions ordered top to bottom; residue positions (PDB 4A0T) shown left to right. **(C-D)** Efficiency of Plating (EOP, left side) and violin plots of plaque area (right side) for (C) *E. coli* UTI1 and (D) *E. coli* UTI2 comparing wild type (WT, grey) and selected variant from the microgravity pool for that strain (right, purple). EOP Data shown as mean ± SD from three biological replicates, normalized to *E. coli* BL21. Significance vs. WT shown as *** (p<0.001).

While variants that performed worse than wildtype (*F_N_*< 0) tended to perform similarly between microgravity and terrestrial conditions (Supplementary Figure 8B), enriched variants (*F_N_* > 0) were highly divergent with no correlation between conditions (Supplementary Figure 8C). Variants enriched in microgravity frequently contained methionine and isoleucine substitutions at interior positions facing the phage (Figure 5A and Supplementary Figure 8D), in contrast to our previous terrestrial results on this host [12]. Substitutions in these areas could influence the tip domain structure to facilitate adsorption with the host receptor in microgravity.

Under terrestrial conditions, top-scoring variants included positively charged substitutions facing the host, consistent with our previous findings on *E. coli* BL21. Additional enriched variants featured negatively charged substitutions (e.g. Q488E, G521D) and glycine substitutions (e.g., G480W, G522P) that may induce structural changes in the tip domain (Figure 5B and Supplementary Figure 8B). These variants were enriched only after prolonged incubation with *E. coli* BL21, suggesting that such substitutions may contribute to long-term infectivity on stationary-phase hosts—an effect not observed in shorter, nutrient-rich conditions.

Because variants enriched in microgravity were highly distinct from those identified under terrestrial conditions—both in this study and in our previous work—we next evaluated whether these substitutions could enhance phage activity terrestrially. If successful, these substitution patterns could be used to improve phage performance without exhaustively sampling the full combinatorial space of the gene. We constructed two combinatorial libraries, each comprising all possible combinations of thirteen top-performing substitutions identified in microgravity (L490I, N502E, F506M, F506Y, F507V, F507Y, P511M, I514M, N531Q, L533K, L533M, A539M, N546I) or under terrestrial conditions (G521H, Q488A, Q488E, G521K, G522P, A547S, G521D, G521E, N502S, I495L, R542H, L533T, F506S). This strategy reduced a potential search space of over 10²¹ variants to fewer than 5,000 per library. Variants were synthesized in an oligo pool, assembled into an unbiased phage library using ORACLE, and passaged terrestrially on two clinically isolated *E. coli* strains (UTI1 and UTI2) that are resistant to wild-type T7 and are associated with urinary tract infections.

We evaluated these pools in efficiency of plating (EOP) experiments and compared their plaquing capability versus wildtype. The combinatorial pool from microgravity showed significant improvement in plaquing efficiency compared to wildtype and had substantially larger plaques, indicating the pool contained variants capable of significantly improving activity on these hosts (Supplementary Figure 9A-B). The terrestrial library performed significantly worse or no better than wildtype. To confirm these results, we isolated individual plaques from the microgravity pool. From UTI1, we recovered a five-substitution variant (L490I, N502E, F507V, L533K, A539M; Variant 1), and from UTI2, a six-substitution variant (L490I, N502E, P511M, L533M, A539M, N546I; Variant 2). These variants demonstrated significantly higher EOP and produced significantly larger plaques on both UTI strains (Figure 5C–D). These findings support our hypothesis that microgravity-enriched substitutions can improve phage performance on terrestrial hosts. The extended incubation in microgravity revealed new mutational hotspots, enabling efficient navigation of sequence space to identify complex variant combinations with enhanced infectivity.

## Discussion

Phage–bacteria interactions play a critical role in shaping microbial ecosystems but remain poorly understood in microgravity. In space, altered collision dynamics and bacterial physiological changes disrupt typical phage–host interplay. Characterizing these interactions provides insight into microbial adaptation in space and reveals novel genes and mechanisms with potential applications on Earth.

We found that phage replication in microgravity was significantly delayed— occurring sometime after the 4-hour time point—but ultimately successful by 23 days, indicating a markedly slower yet productive replication cycle. Numerous significantly enriched *de novo* mutations were identified in both phage and bacterial genes under microgravity and terrestrial conditions, suggesting strong selective pressures in both environments. In microgravity, structural genes *gp7.3*, *gp11*, and *gp12* emerged as particularly important, while enrichment of mutations in the non-structural gene *gp0.5* suggests its putative association with the host membrane may contribute more to phage fitness than previously recognized [22]. The overall distribution of mutations highlights genomic regions that warrant further investigation in future studies.

Mutations in the bacterial host were predominantly found in genes involved in outer membrane structure, stress response, and nutrient management. These findings are consistent with previous studies showing upregulation of similar gene classes in closely related *E. coli* strains under microgravity conditions. Such genes play key roles in managing environmental stress, regulating nutrient availability, and facilitating transmembrane transport in this unique setting [11,49–52]. The high frequency of mutations in these genes suggests they may reduce phage infectivity, offering bacteria an additional selective advantage.

Results from the T7 receptor binding protein (RBP) tip domain DMS library revealed significantly different selection patterns in microgravity compared to terrestrial incubation, both in this study and in our previous work [12]. Microgravity-enriched trends enabled efficient navigation of sequence space, leading to multi-substitution variants with significantly enhanced activity against uropathogenic *E. coli* under terrestrial conditions. Notably, combinatorial variants derived from terrestrial-enriched mutations failed to outperform wild-type, suggesting that the unique selective pressures of microgravity uncovered previously unrecognized functional regions with terrestrial relevance.

This study focused on a single non-motile strain of *E. coli*. Motile elements could potentially enhance fluid mixing, and future studies incorporating a broader range of bacterial strains would help clarify this effect. Additionally, the experimental design included several freeze–thaw cycles and a delay in processing, which likely reduced phage and bacterial viability. While some of these limitations are inherent to space-based research, minimizing freeze–thaw events and processing delays in future experiments could improve sample integrity and data quality.

Our study offers a preliminary look at how microgravity influences phage–host interactions. Exploring phage activity in non-terrestrial environments reveals novel genetic determinants of fitness and opens new avenues for engineering phages for terrestrial use. The success of this approach helps lays the groundwork for future phage research aboard the ISS.

## Supplementary Files

**Supplementary File 1** – Summary of Cryotube Biocompatibility Testing, Cryotube Freeze-Thaw Testing and Experiment Verification Testing

**Supplementary File 2** – Summary of *de novo* Bacteriophage Mutations

**Supplementary File 3** – Summary of *de novo* Bacterial Mutations

**Supplementary File 4** – Summary of Deep Mutational Scanning Results

**Supplementary Figure 1.**
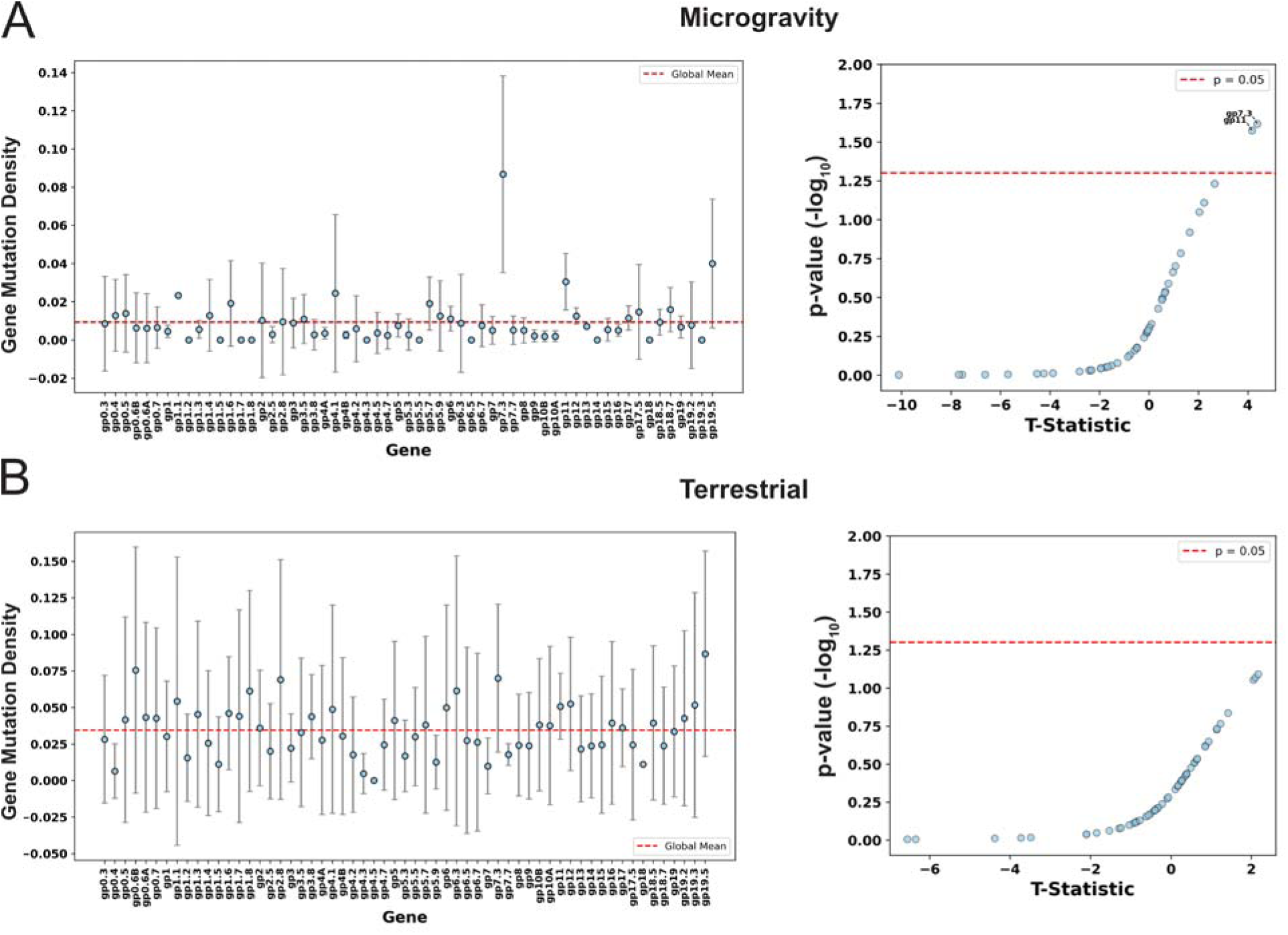
One-tailed 95% confidence intervals for mutation density (non-synonymous substitutions and frameshift count divided by protein length) for each phage gene after incubation in **(A)** microgravity and **(B)** after terrestrial incubation. Right panels show t-statistics and corresponding p-values from one-tailed t-tests with FDR correction. Genes with adjusted p-values <0.05 were considered significant.

**Supplementary Figure 2.**
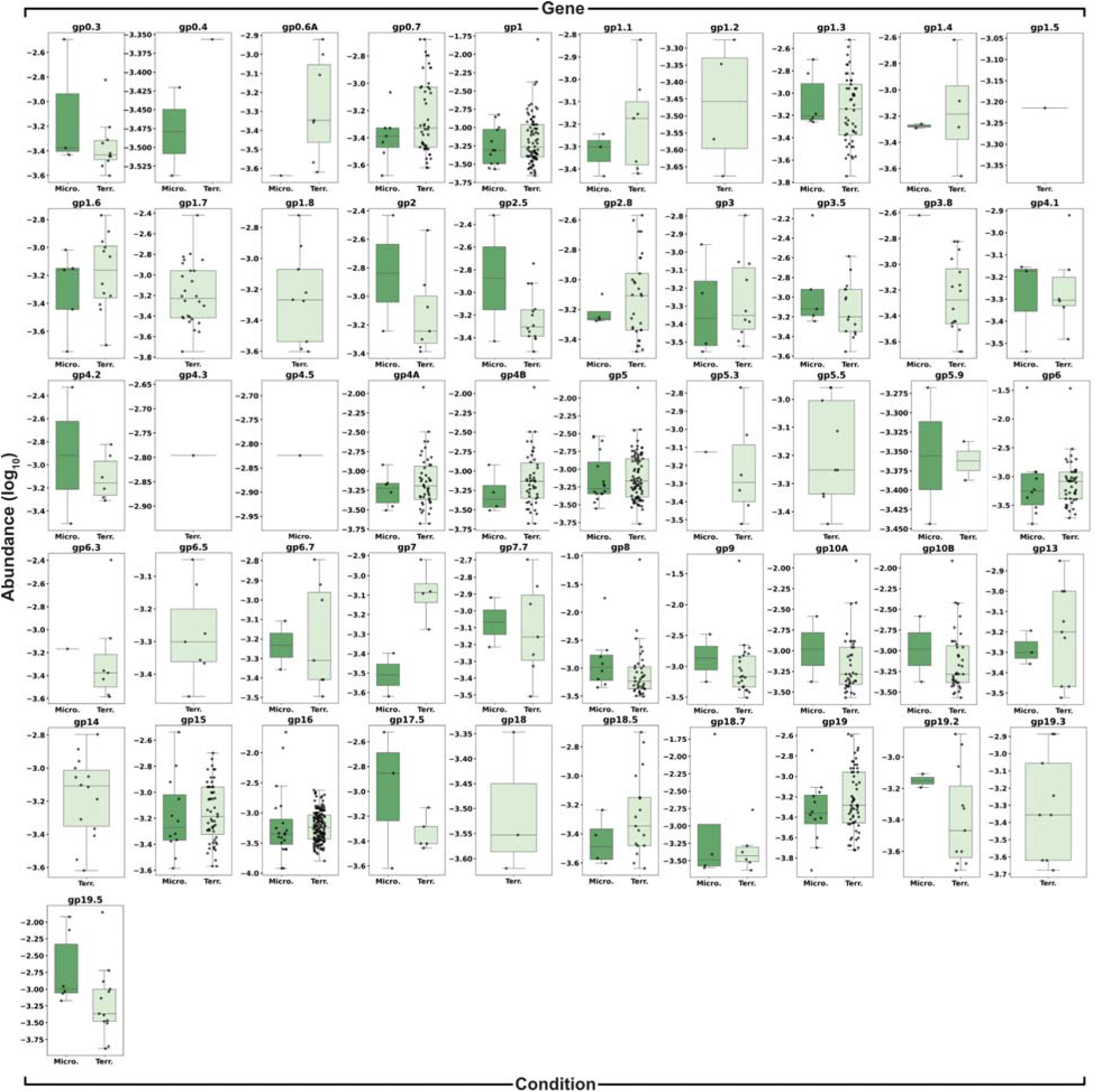
Comparison of log_10_ abundance of *de novo* non-synonymous substitutions and frameshifts for phage genes not shown in Figure 2 after incubation in microgravity (left, dark green, Micro.) and terrestrially (right, light green, Terr.). No significant differences were detected, or data were too sparse to assess significance. Genes with mutations in only one condition show only that condition.

**Supplementary Figure 3.**
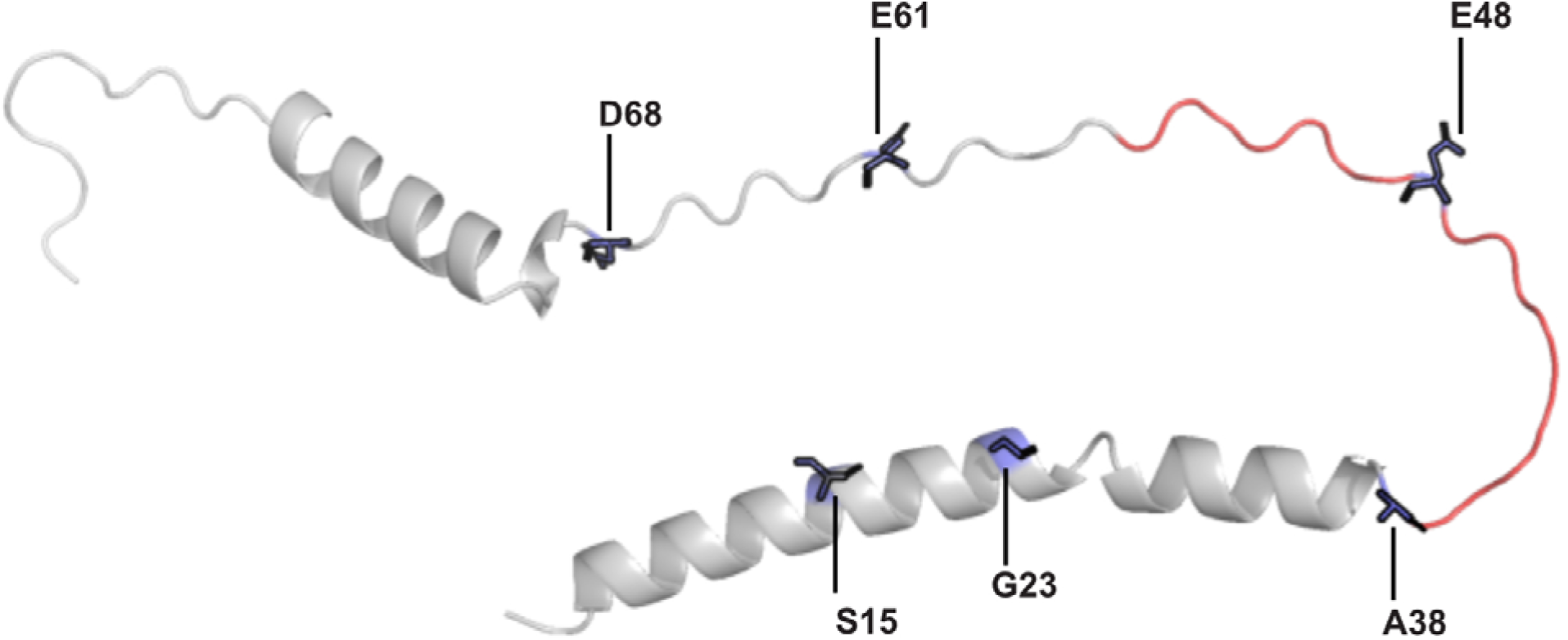
Predicted structure of *gp7.3* using Alphafold2. Positions with significantly enriched substitutions in microgravity are shown in blue. The unstructured region from G39 to Q55 related to in-frame deletions is shown in red.

**Supplementary Figure 4.**
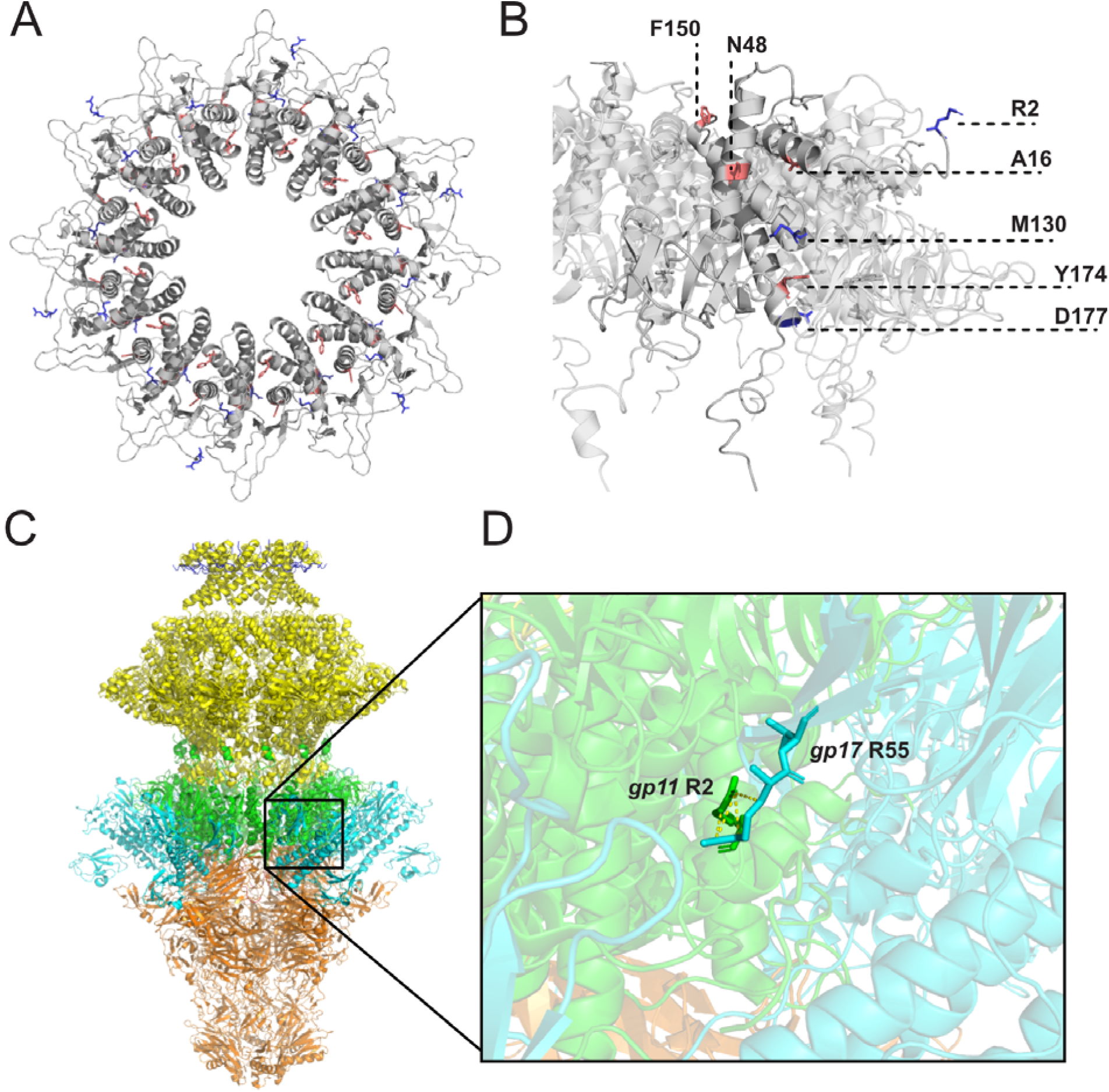
**(A)** Electron microscopy structure of the *gp11* complex (PDB 7BOX) showing all 12 subunits. Mutations significantly enriched in microgravity are shown in blue; those enriched terrestrially in red. **(B)** Enlarged view of *gp11* with one subunit labeled, highlighting enriched mutations using the color scheme from (A). **(C)** Electron microscopy structure of the T7 portal-tail complex (PDB 9JYZ). *gp12* (portal protein) is shown in orange, *gp11* in green, *gp17* attachment in blue, and *gp8* (extending toward core proteins) in yellow. **(D)** Close-up of the interaction between R2 in *gp11* (green) and *R55* in gp17 (blue), with yellow dashed lines indicating contacts within 3.5 Å. R2 resides in an unstructured, flexible region that may interact with multiple *gp17* residues.

**Supplementary Figure 5.**
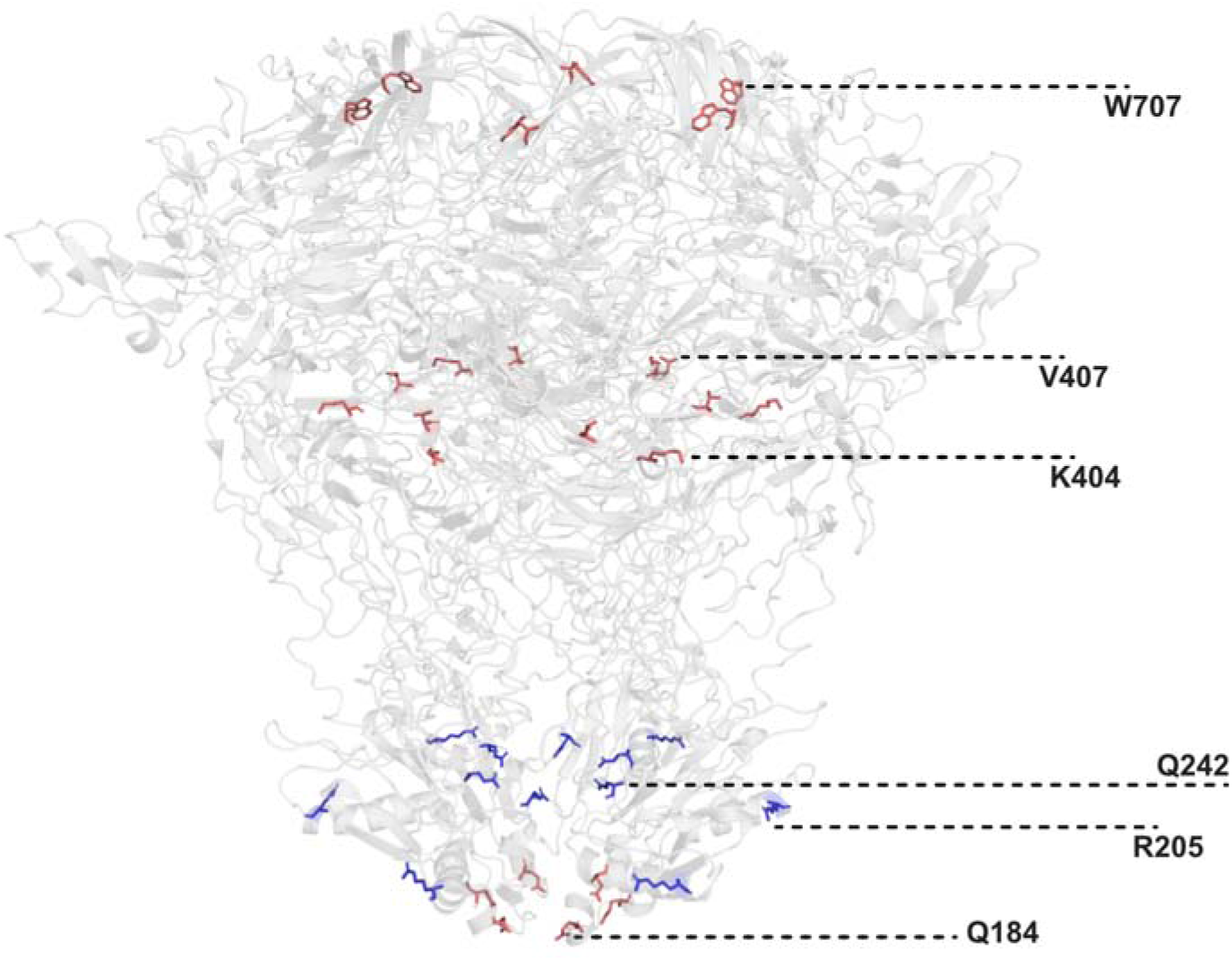
**(A)** Electron microscopy structure of the *gp12* complex (PDB 7BOY) showing all six subunits. Mutations significantly enriched in microgravity are shown in blue; those significantly enriched terrestrially in red. Positions are labeled on one representative subunit.

**Supplementary Figure 6.**
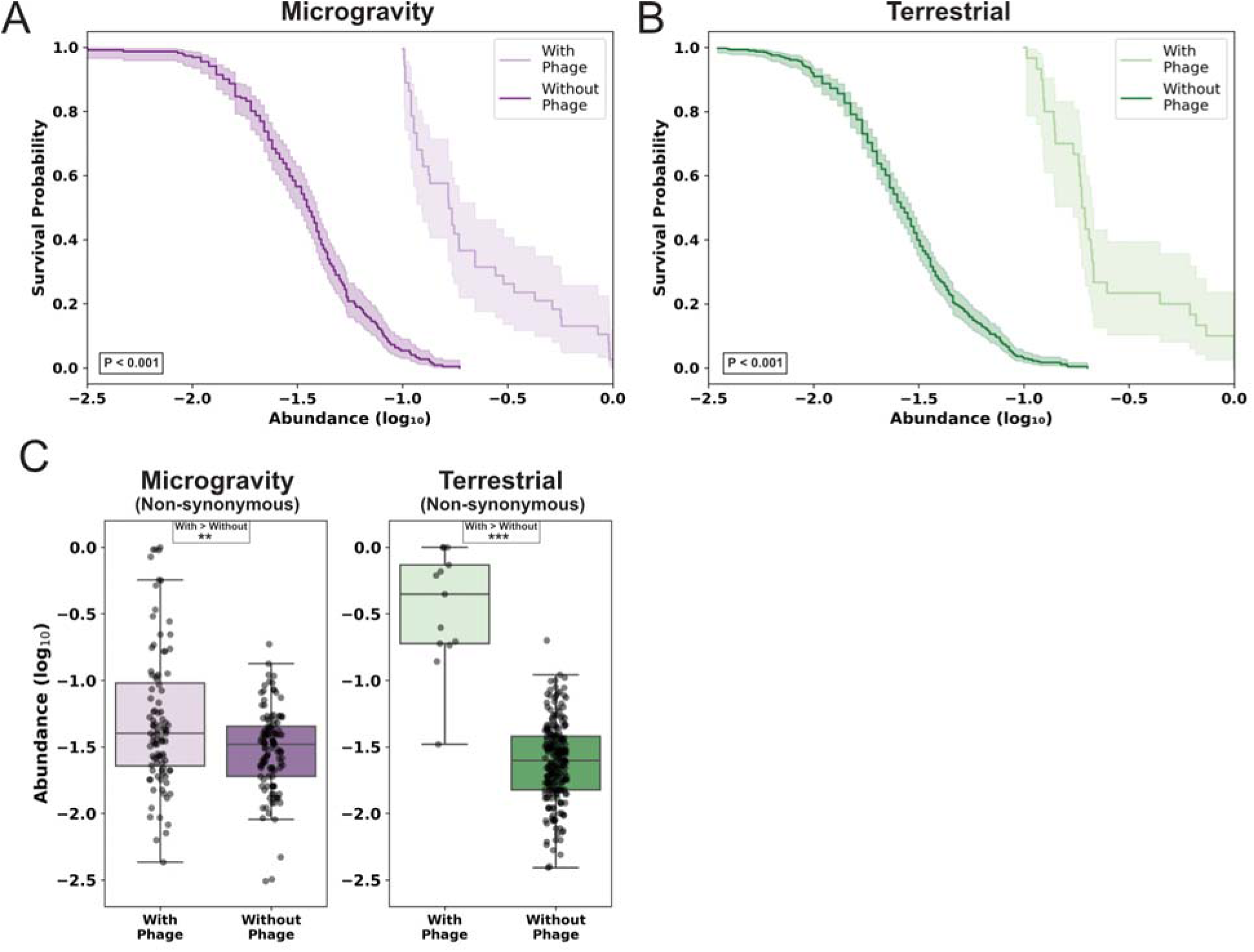
**(A-B)** Kaplan–Meier survival curves for bacterial non-synonymous substitutions and frameshift mutations after incubation (A) terrestrially or (B) in microgravity, with phage (light purple/light green) or without phage (dark purple/dark green). Shaded regions represent 95% confidence intervals. Survival probability reflects the proportion of mutations with abundance above the averaged limit of detection for each condition (log_10_ −1 with phage, log_10_ −2.5 without). P-values were calculated using log-rank tests. **(C)** Boxplots of log_10_ abundance for non-synonymous substitutions and frameshift mutations after microgravity (left) or terrestrial (right) incubation, comparing conditions with (light shading) and without (dark shading) phage. Significance was assessed using a Mann–Whitney U test (*p < 0.05, **p < 0.01, ***p < 0.001), with “>” indicating the more abundant group.

**Supplementary Figure 7.**
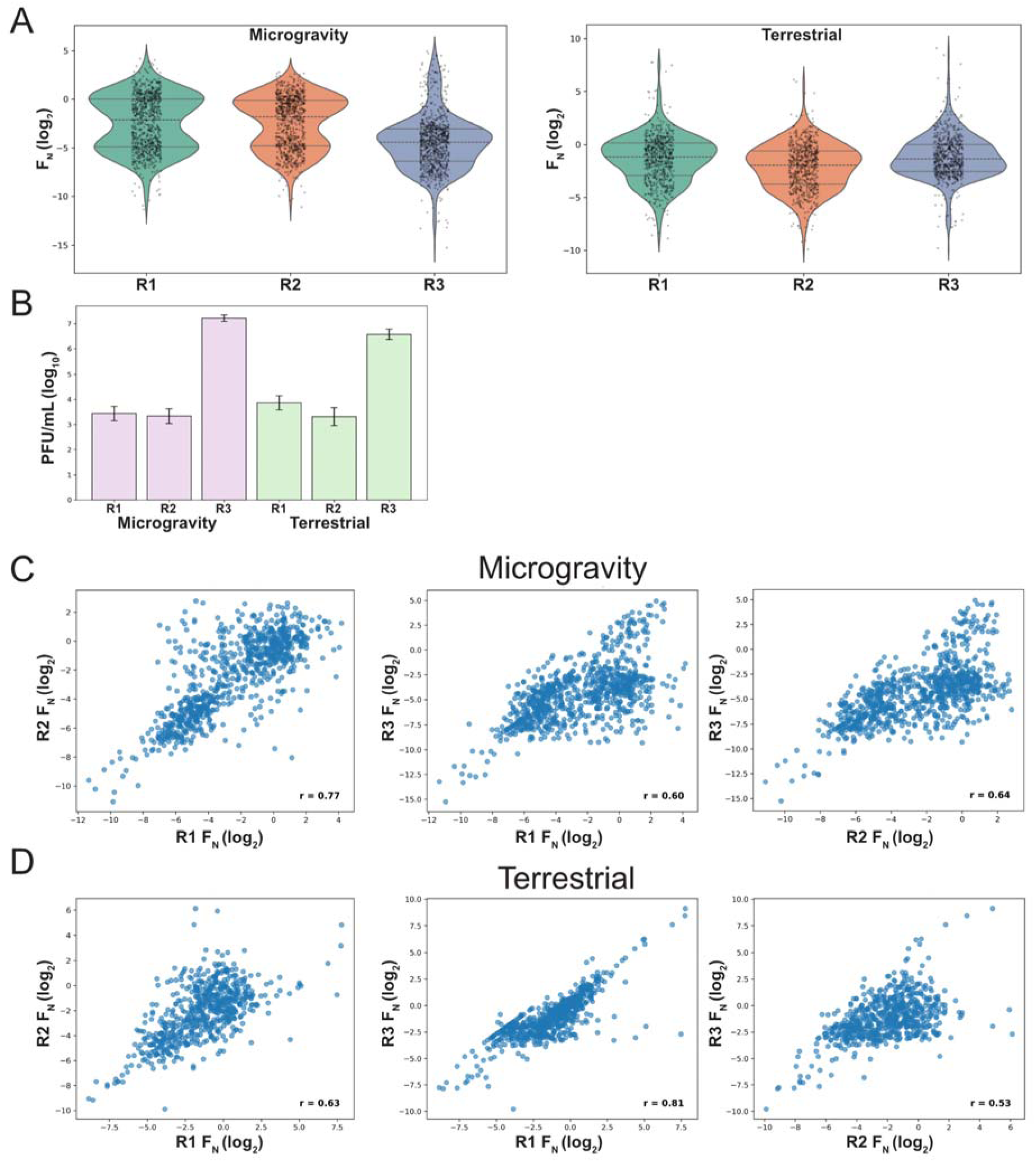
**(A)** Violin plots of DMS variant’s F_N_ (log_2_) scores across biological replicates (R1-3) in microgravity (left) and terrestrial (right) conditions. **(B)** Phage titer (log_10_ PFU) for DMS replicates 1, 2, and 3 (R1, R2, and R3) after microgravity (left, purple) and terrestrial (right, green) incubation, shown as mean ± SD. **(C-D)** Correlation plot of DMS variants F_N_ (log_2_) score between replicates (R1-3) in (C) microgravity and (D) terrestrial conditions. Pearson’s r is displayed on the bottom right of each plot.

**Supplementary Figure 8.**
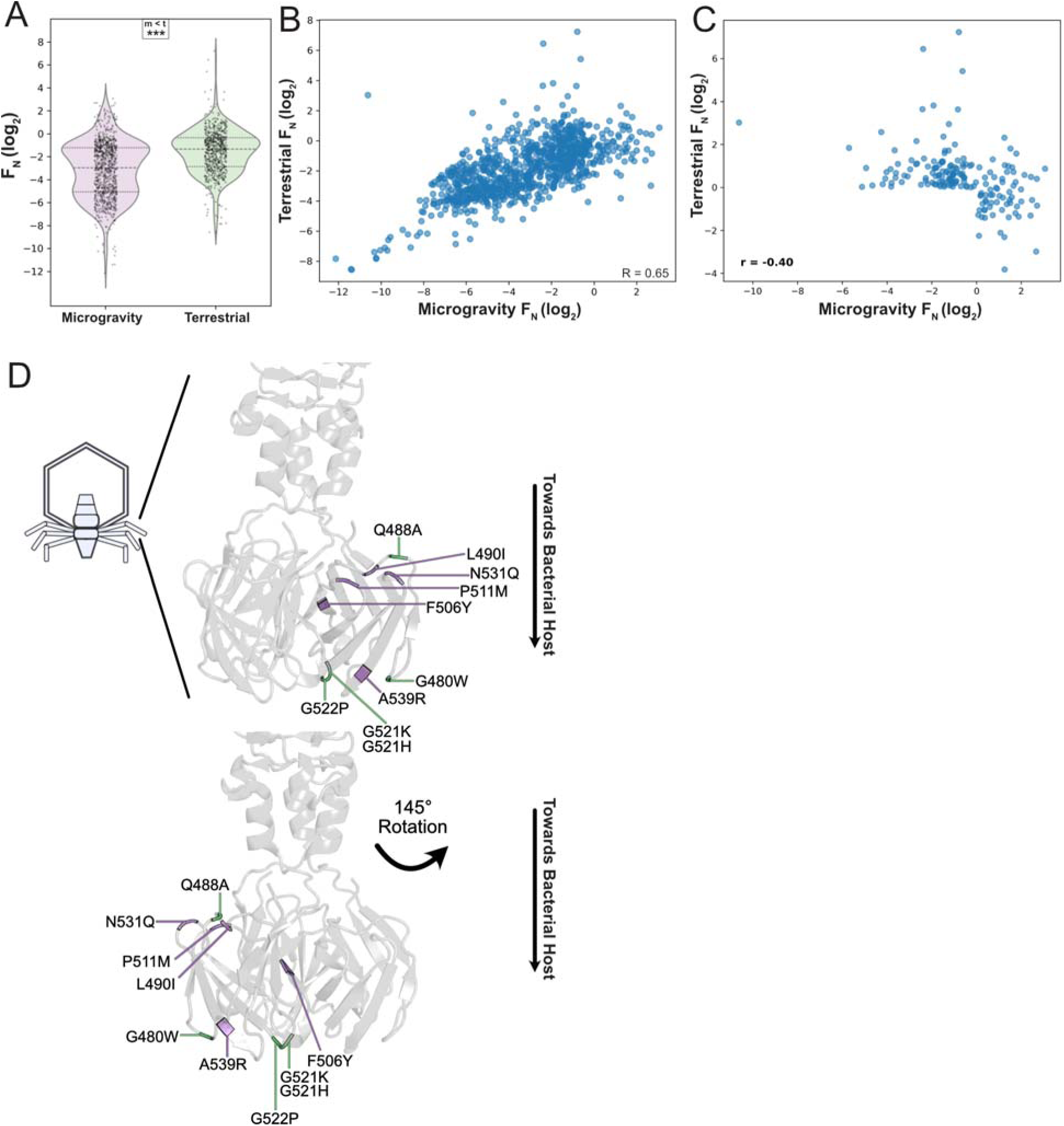
**(A)** Violin plots of average *F_N_* (log_2_) for DMS variants in microgravity (left, purple) and terrestrial (right, green) conditions. Significance indicated as *** (p<0.001). **(B)** Correlation plot of F_N_ (log_2_) scores for all DMS variants between microgravity and terrestrial conditions. Pearson’s *r* displayed in the bottom right. **(C)** Correlation of enriched (*F_N_* (log_2_) > 0) variants between conditions. Pearson’s r shown bottom left. **(D)** Crystal structure and secondary structure topology of the RBP tip domain (PDB: 4A0T), with substitutions enriched in microgravity (purple) or terrestrial (green) conditions highlighted. Two views are shown for clarity with a 145° rotation.

**Supplementary Figure 9.**
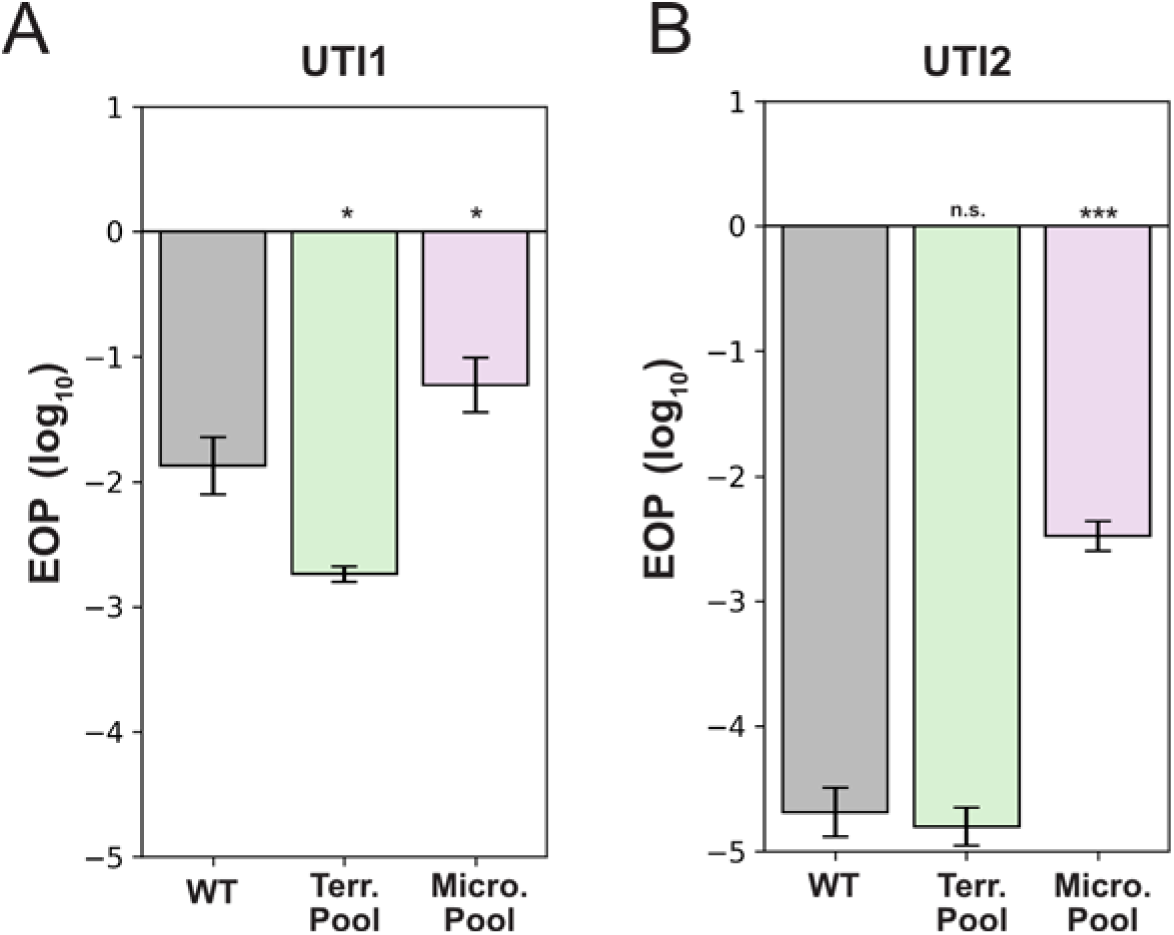
**(A-B)** Efficiency of Plating (EOP) results on (A) *E. coli* UTI1 and (B) *E. coli* UTI2 comparing wild type (WT, left, grey), terrestrial combinatorial pool (middle, green), and microgravity combinatorial pool (right, purple). Data shown as mean ± SD from three biological replicates, normalized to *E. coli* BL21. Significance vs. WT shown as * (p< 0.05), *** (p<0.001), or n.s. (not significant).

## Contributions

P.H.: Conceptualization, Data curation, Software, Formal analysis, Investigation, Visualization, Methodology, Writing - original draft, Writing - review and editing

C.C.: Conceptualization, Data curation, Formal analysis, Investigation, Methodology, Writing - original draft, Writing - review and editing

A.M.: Software

K.N.: Software

H.M., R.O., O.H.: Resources, Project administration, Methodology, Writing - review and editing

S.R.: Conceptualization, Resources, Supervision, Funding acquisition, Project administration, Writing - review and editing

## Acknowledgements

We thank Dr. R. Welch for UTI strains. This work was supported by the Defense Threat Reduction Agency (Grant HDTRA1-16-1-0049). C.C was supported by a graduate training scholarship from the Anandamahidol Foundation (Thailand).

## Methods

### Phage and Bacterial Strains

*T7 bacteriophage* was obtained from ATCC (ATCC® BAA-1025-B2). The T7 deep mutational scanning (DMS) library used in this study was the same library stock generated in our previous work [12]. T7 acceptor phages used for ORACLE-based construction of the combinatorial libraries were also created as previously described [12]. *Escherichia coli BL21* was sourced from laboratory stocks. Uropathogenic *E. coli* strains UTI1 and UTI2 were provided by Dr. R. Welch (University of Wisconsin, Madison) and originate from a urinary tract infection isolate collection [53].

T7 phage was initially propagated on *E. coli* BL21 following receipt from ATCC and subsequently on appropriate hosts as described in specific experimental sections. All phage experiments were performed using LB media and the same culture conditions used for bacterial hosts. Phages were stored in LB at 4°C for short-term use. For long-term storage, microbial samples were frozen at –80°C in 100% LB media.

### Media and Culture Conditions

All bacterial strains were cultured in Luria-Bertani (LB) media consisting of 1% tryptone, 0.5% yeast extract, and 1% NaCl in deionized water. LB plates were supplemented with 1.5% agar, while top agar used for phage plating contained 0.5% agar. LB media was used for all experiments, including bacterial recovery and phage propagation. All incubations were carried out at 37°C without shaking, in either terrestrial or microgravity environments as appropriate.

### Sample Preparation and Handling

Phage and bacterial stock titer were confirmed and samples were prepared by mixing 4 mL of *E. coli* BL21 in exponential phase (∼1 × 10^8^ CFU/mL) with the appropriate amount of T7 phages in Rhodium Cryotubes. Samples were immediately frozen at –80°C and shipped to NASA as described.

The initial planned time points for incubation were 1, 2, and 3 hours, and 25 days; however, actual time points were adjusted on the ISS to accommodate astronaut scheduling. Final incubation time points were 1, 2, and 4 hours, and 23 days. The duration of incubation aboard the ISS was recorded precisely, and terrestrial control samples were incubated for matching durations, based on the actual timepoints rather than the proposed schedule. This approach was necessary because real time tracking of the samples was not possible, so microgravity and terrestrial samples could not be incubated in parallel accurately. Terrestrial samples are thus frozen for a longer duration than microgravity samples. After incubation samples were refrozen, shipped to our laboratory, and then thawed at 37°C and immediately split for genomic DNA extraction, PCR for DMS, and titering of both phage and bacteria.

### Titering Phage

For samples returned for processing, 1 mL of each sample was centrifuged at 16g for 1 minute, and the supernatant was filtered through a 0.22 μm filter. To determine phage titer, titer was first estimated by spot plates and then confirmed by whole plate efficiency of plating (EOP) assays. Samples were serially diluted (1:10 or 1:100) in LB to a final dilution of up to 10^-8^ in 1.5 mL microcentrifuge tubes. Spot assays were performed by mixing 250 μL of stationary-phase bacterial host with 3.5 mL of 0.5% top agar. The mixture was briefly vortexed and plated onto LB agar plates pre-warmed to 37°C. Once the top agar solidified (∼5 minutes), 1.5 μL of each phage dilution was spotted onto the plate in series. Plates were incubated at 37°C and checked after 20–30 hours to estimate titer. Titers were then confirmed via full-plate plaque assays.

For whole-plate EOP assays, 400 μL of exponentially growing bacterial culture was mixed with 5–50 μL of diluted phage, aiming to achieve ∼50 plaque-forming units (PFUs) per plate after overnight incubation. The phage–host mixture was briefly vortexed and centrifuged, then combined with 3.5 mL of 0.5% top agar. After a brief vortex, the mixture was immediately poured onto LB plates pre-warmed to 37°C. Plates were allowed to solidify (∼5 minutes), inverted, and incubated overnight. PFUs were counted after 20–30 hours, and final phage titers were calculated from these counts.

### Titering Bacteria

Bacterial concentrations were determined via serial dilution (1:10 or 1:100 in LB) and plating. From each dilution, 100 μL was plated and spread using sterile beads to target ∼50 colony-forming units (CFUs) per plate. Plates were incubated overnight at 37°C and counted the following day. For *E. coli* BL21, three independent dilution series were performed to correlate OD_600_ values with CFU/mL and ensure accurate bacterial concentrations during phage mixing for experimental sample preparation.

### PCR and Sequencing

All PCR reactions were performed using KAPA HiFi DNA Polymerase (Roche KK2101). The combinatorial library was generated using the ORACLE method, as previously described [12]. Cloning procedures followed manufacturer instructions unless otherwise specified.

For whole-genome sequencing (WGS), phage genomes were extracted using the Norgen Biotek Phage DNA Isolation Kit (Cat. 46800), and bacterial genomic DNA was extracted using the Norgen Biotek Bacterial Genomic DNA Isolation Kit (Cat. 17900). Genomic DNA libraries were prepared using the Illumina DNA Prep kit (Cat. 20060060) and sequenced on an Illumina NextSeq 1000 platform.

PCR reactions for amplification of the DMS and combinatorial libraries used 1 μL of undiluted phage lysate directly as template (DNA isolation is not required), with an extended denaturation step of 5 minutes at 95°C. For low phage titers in DMS samples, PCR and next-generation sequencing (NGS) failed using this approach, presumably because of reduced template in these samples. To overcome this, we concentrated the all of the remaining volume of each sample (∼2 mL) approximately 100-fold using Pierce™ Protein Concentrators PES, 10K MWCO (Cat. 88513) and used 3 μL of the concentrated sample per PCR reaction to enabling successful amplification and analysis. For plaque analysis on UTI strains, small plaque samples were picked directly and used as PCR template. Detailed cloning protocols are available upon request.

### General Data Analysis

Multiplicity of infection (MOI) was calculated by dividing the phage titer by the corresponding bacterial concentration. The MOI for the T7 DMS library was estimated using a helper plasmid, as described previously [12].

Efficiency of Plating (EOP) values were calculated using *E. coli* BL21 as a reference host. EOP was defined as the phage titer on the test host divided by the titer on the reference host, followed by log_10_ transformation. Values are reported as mean ± standard deviation (SD).

Deep sequencing was performed to evaluate phage populations as described previously [12]. Phage sequencing achieved an average depth of ∼49,000× per base across the genome, enabling detection of low-abundance mutations. Bacterial sequencing depth averaged ∼250× per base in phage-mixed samples and ∼1,300× in phage-free samples, limiting mutation analysis in the former to more abundant variants.

Whole-genome sequencing (WGS) mutations were identified using Breseq [54]. For Figure 3, genes were grouped based on Gene Ontology (GO) classifications [23,24]:

- Membrane-associated genes: GO:0016020 (Membrane), GO:0009103 (LPS biosynthesis), GO:0030288 (Outer membrane bound periplasmic space), GO:0042597 (Periplasmic space).
- Metabolism-associated genes: GO:0008152 (Metabolic process), GO:0019222 (Regulation of metabolic process).

### Statistical Analysis

To evaluate whether non-synonymous *de novo* substitutions and frameshift mutations were significantly enriched compared to synonymous substitutions, we compared the frequency of each non-synonymous substitutions and frameshift (phage: >1% abundance; bacteria: >25% abundance) to the distribution of synonymous mutations using a one-sided Mann-Whitney U test with Benjamini-Hochberg false discovery rate (FDR) correction (scipy.stats.mannwhitneyu, statsmodels.stats.multitest.multipletests, method = ‘fdr_bh’). Adjusted p-values < 0.05 were considered significant. This approach assumes that after 23 days of selection the distribution of synonymous substitutions approximates either a neutral baseline or reflects minimal selective pressure, with the benefit that if there is positive selection for synonymous substitutions there would be no increase in false positives using this method.

To determine whether non-synonymous *de novo* substitutions and frameshift mutations were more abundant in bacterial samples exposed to phage, we applied the Mann-Whitney U test (scipy.stats.mannwhitneyu) to compare mutation frequencies across groups [55]. Due to high detection limits in phage-mixed samples, we also performed left-censored data analysis using Kaplan-Meier survival curves (lifelines.KaplanMeierFitter) and applied a log-rank test (lifelines.statistics.logrank_test) to assess significant differences in mutation distributions between groups [56,57].

Mutation density in phage genes was calculated by dividing the number of non-synonymous *de novo* substitutions and frameshift mutations by the length (in amino acids) of each protein product. To assess whether any gene had significantly higher mutation density, we compared individual gene densities to the condition-specific average using a one-tailed t-test with Benjamini-Hochberg FDR correction (scipy.stats.ttest_1samp, statsmodels.stats.multitest.multipletests, method = ‘fdr_bh’, alternative = ‘greater’). Additionally, one-tailed 95% confidence intervals were calculated using scipy.stats.t.ppf and visualized in volcano plots.

### Structural Visualization

Structural model images were generated using the PyMOL Molecular Graphics System, Version 3.0 (Schrödinger, LLC). *Gp7.3* structure was predicted using AlphaFold2 and ColabFold with MMseqs2, using the predicted structure with the highest confidence [58–60]. Electron Microscopy images were based on PDB 7BOX (gp11) and PDB 7BOY (gp12) [20]. A composite structure of the T7 portal–tail complex was generated using PDB 9JYZ, from an upcoming publication. Numbering for DMS and combinatorial library positions are based on PDB 4A0T [43].

## Notes

### Competing Interest Statement

The authors have declared no competing interest.

### Summary of Updates

Manuscript has been updated with additional analysis and terrestrial experiments.

