## Supplementary File 1 for "Bacteriophage-host interactions in microgravity aboard the International Space Station"

**RHODIUM SCIENTIFIC, LLC**
1300 BAY AREA BLVD, STE B275-5
HOUSTON, TX 77058

MAIN: 210.641.2880

### STANDARD OPERATING PROCEDURE

### RHODIUM CRYOTUBE BIOCOMPATIBILITY TEST PROTOCOL

**SOP-BIOCOMPAT-004**

**Rev 0**

PREPARED BY: Dr. Heath Mills, Chief Scientific Officer

REVIWED BY: Olivia Holzhaus, Chief Executive Officer

DATE: 08-18-19

This document does not contain technical data as defined in the ITAR, 22 CFR 120.10; or technology as defined under the EAR (15 CFR 730-774).

RHODIUM SCIENTIFIC, LLC
1300 BAY AREA BLVD, STE B275-5
HOUSTON, TX 77058

MAIN: 210.641.2880
WWW.RHODIUMSCIENTIFIC.COM

#### REVISION HISTORY

| REV | CHANGE | SECTION AFFECTED |
|---|---|---|
| 0 | Initial Release, 08-18-19 | All |

### RHODIUM CRYOTUBE BIOCOMPATIBILITY TEST PROTOCOL

**Preformed by:**
**Test POC** – Dr. Srivatsan Raman, Assistant Professor, University of Wisconsin-Madison
Phillip Huss, Graduate Research Assistant, University of Wisconsin-Madison

**Directed by:**
**Test POC** – Dr. Heath J. Mills, CSO, Rhodium Scientific, LLC
Dr. R.P. Oates, Program Manager, Rhodium Scientific, LLC
Olivia Holzhaus, CEO, Rhodium Scientific, LLC

**Test Purpose:**
Determine biological and chemical effects to Rhodium Cryotube integrity following incubation.

**Test Configuration:**
A total of twelve (12) Rhodium Cryotubes (Part Number: RhCT-0005) were filled with 4.0 mL of LB growth media inoculated with *E. coli* and bacteriophage. This is the same media, bacteria and bacteriophage that will be used during the Phage Evolution experiment. This test configuration matches the flight configuration.

**Test Protocol:**
1. Pipette 3.8 mL LB media into twenty four (24) Rhodium Cryotubes labeled #1-12 EC and #1-#12 ECP. All cryotubes #1-3 will be additionally labeled "1 hr", cryotubes #4-6 labeled "2 hr", cryotubes #7-9 labeled "4 hr", and cryotubes #10-12 labeled "25 days".
2. Pipette 0.150 mL (150 µL) *E. coli* from overnight culture into all Rhodium Cryotubes.
3. Pipette 0.05 mL (50 µL) of bacteriophage into cryotubes labeled ECP. Pipette 0.05 mL (50 µL) LB media into cryotubes labeled EC.
4. Record the weight of all labeled Rhodium Cryotubes in Table 1.
5. Place loaded Rhodium Cryotubes in the -80$^o$C freezer for 30 minutes.
6. Remove the Rhodium Cryotubes from the freezer and transfer directly to an incubator preconditioned to 37$^o$C.
7. Incubate the cryotube sets according to their designated incubation times of 1 hr, 2 hr, 4 hr and 37 days.
8. At the end of each incubation period, remove the respective samples from the incubator and observe each cryotube's structural integrity by pressing firmly against the tube. Verify there is no cracking, leaking or deterioration to the tubes.
9. Visually inspect the volume within each tube to qualitatively determine loss of liquid. Record the visual volume in Table 1.
10. After the visual volume has been recorded, re-weigh and record the weight of each cryotube to determine variances from initial weights and to quantitatively assess any loss of fluid.
11. Return cryotubes to freezer at -80°C.

**Table 1: Biocompatibility Test Results – Bacteria only**

| Column A | Column B | Column C | Column D | Column E | Column F | Column G |
|---|---|---|---|---|---|---|
| **Tube Number** | **Incubation Time (at 37°C)** | **Initial Visual Vol. (mL)** | **Final Visual Vol. (mL)** | **Initial Weight (g)** | **Final Weight (g)** | **Percent Variance (%)** |
| **1 EC** | 1 hr | 4.0 | 4.0 | 7.0854 | 7.0844 | 0.01% |
| **2 EC** | 1 hr | 4.0 | 4.0 | 7.0364 | 7.0358 | 0.01% |
| **3 EC** | 1 hr | 4.0 | 4.0 | 7.0583 | 7.0580 | 0.00% |
| **4 EC** | 2 hr | 4.0 | 4.0 | 7.0643 | 7.0362 | 0.40% |
| **5 EC** | 2 hr | 4.0 | 4.0 | 7.0525 | 7.0522 | 0.00% |
| **6 EC** | 2 hr | 4.0 | 4.0 | 7.0688 | 7.0673 | 0.02% |
| **7 EC** | 4 hr | 4.0 | 4.0 | 7.0651 | 7.0623 | 0.04% |
| **8 EC** | 4 hr | 4.0 | 4.0 | 7.0724 | 7.0597 | 0.18% |
| **9 EC** | 4 hr | 4.0 | 4.0 | 7.0849 | 7.0619 | 0.32% |
| **10 EC** | 37 days | 4.0 | 4.0 | 7.0643 | 7.0273 | 0.52% |
| **11 EC** | 37 days | 4.0 | 4.0 | 7.1000 | 7.0601 | 0.56% |
| **12 EC** | 37 days | 4.0 | 4.0 | 7.0878 | 7.0334 | 0.77% |

**Table 2: Biocompatibility Test Results – Bacteria and Phage**

| Column A | Column B | Column C | Column D | Column E | Column F | Column G |
|---|---|---|---|---|---|---|
| **Tube Number** | **Incubation Time (at 37°C)** | **Initial Visual Vol. (mL)** | **Final Visual Vol. (mL)** | **Initial Weight (g)** | **Final Weight (g)** | **Percent Variance (%)** |
| **1 ECP** | 1 hr | 4.0 | 4.0 | 7.0861 | 7.0857 | 0.01% |
| **2 ECP** | 1 hr | 4.0 | 4.0 | 7.0674 | 7.0672 | 0.00% |
| **3 ECP** | 1 hr | 4.0 | 4.0 | 7.0665 | 7.0665 | 0.00% |
| **4 ECP** | 2 hr | 4.0 | 4.0 | 7.1106 | 7.1104 | 0.00% |
| **5 ECP** | 2 hr | 4.0 | 4.0 | 7.1048 | 7.1039 | 0.01% |
| **6 ECP** | 2 hr | 4.0 | 4.0 | 7.1151 | 7.1146 | 0.01% |
| **7 ECP** | 4 hr | 4.0 | 4.0 | 7.1219 | 7.1208 | 0.02% |
| **8 ECP** | 4 hr | 4.0 | 4.0 | 7.0541 | 7.0533 | 0.01% |
| **9 ECP** | 4 hr | 4.0 | 4.0 | 7.0411 | 7.0408 | 0.00% |
| **10 ECP** | 37 days | 4.0 | 4.0 | 7.0951 | 7.0735 | 0.30% |
| **11 ECP** | 37 days | 4.0 | 4.0 | 7.0736 | 7.0619 | 0.17% |
| **12 ECP** | 37 days | 4.0 | 4.0 | 7.0872 | 7.0724 | 0.21% |

### DEMONSTRATION OF CAPABILITY

#### RHODIUM CRYOTUBE BIOCOMPATIBILITY TEST PROTOCOL

#### SOP-BIOCOMPAT-004

#### Rev 0

| DATE | DEMONSTRATOR | SIGNATURE |
|---|---|---|

**RHODIUM SCIENTIFIC, LLC**
1300 BAY AREA BLVD, STE B275-5
HOUSTON, TX 77058

MAIN: 210.641.2880

### STANDARD OPERATING PROCEDURE

### RHODIUM CRYOTUBE FREEZE-THAW TEST PROTOCOL

#### SOP-FRETHAW-003
#### Rev 0

PREPARED BY: Dr. Heath Mills, Chief Scientific Officer

REVIWED BY: Olivia Holzhaus, Chief Executive Officer

DATE: 08-18-19

This document does not contain technical data as defined in the ITAR, 22 CFR 120.10; or technology as defined under the EAR (15 CFR 730-774).

| REVISION HISTORY | | |
|---|---|---|
| **REV** | **CHANGE** | **SECTION AFFECTED** |
| 0 | Initial Release, 08-18-19 | All |

RHODIUM SCIENTIFIC, LLC
1300 BAY AREA BLVD, STE B275-5
HOUSTON, TX 77058

MAIN: 210.641.2880

### RHODIUM CRYOTUBE FREEZE/THAW TEST PROTOCOL

**Preformed by:**
**Test POC** – Dr. Srivatsan Raman, Assistant Professor, University of Wisconsin-Madison
Phillip Huss, Graduate Research Assistant, University of Wisconsin-Madison

**Directed by:**
**Test POC** – Dr. Heath J. Mills, CSO, Rhodium Scientific, LLC
Dr. R.P. Oates, Program Manager, Rhodium Scientific, LLC
Olivia Holzhaus, CEO, Rhodium Scientific, LLC

**Test Configuration:**
A total of eighteen (18) Rhodium Cryotubes were filled with 4.0 mL of sterile LB growth media. This is the same media that will be used during the proposed Phage Evolution experiment. This configuration matches the flight configuration.

**Test Protocol:**
1. Pipette 4.0 mL LB media into eighteen (18) individual Rhodium Cryotubes labeled 1-18. Cryotubes #1-6 should be labeled with “5 min” for short duration tests, cryotubes #7-12 should be labeled with “20 days” for medium duration tests, and cryotubes #13-18 should be labeled “75 days” for long duration tests.
2. In Table 1, record the initial visual volume (Column C) and the initial weight (Column E) of all Rhodium Cryotubes
3. Store Rhodium Cryotubes #7-18 in a -80°C freezer for the medium and long duration tests.
4. Place Rhodium Cryotubes #1-6 in an ethanol/liquid nitrogen solution (~196$^{o}$C) for 5 minutes.
5. After the 5 minutes, thaw tubes #1-6 by placing in an incubator preconditioned to 37$^{o}$C for 30 minutes.
6. After incubation for 30 minutes, place cryotubes #1-6 back in the ethanol/liquid nitrogen solution (~196$^{o}$C) for an additional 5 minutes,
7. Thaw cryotubes in the preconditioned incubator at 37°C for 30 minutes.
8. After the 30 minute incubation, remove the cryotubes, observe each cryotube for structural integrity by pressing firmly against the tube and for cracks, leaks, or deterioration in the tube.
9. In Table 1, record the cryotubes final volume visually (Column D) and the final weight (Column F) to qualitatively determine any loss of liquid.
10. After 37 days, remove cryotubes #7-12 from -80°C freezer and repeat steps 7-9 for the medium duration Freeze-Thaw Test.
11. After a minimum of 75 days, remove cryotubes #13-18 from -80°C freezer and repeat steps 7-9 for the long duration Freeze-Thaw Test.

**Table 1: Freeze-Thaw Test Results**

| Column A | Column B | Column C | Column D | Column E | Column F | Column G |
|---|---|---|---|---|---|---|
| **Tube Label Number** | **Duration of Time Frozen** | **Initial Visual Vol. (mL)** | **Final Visual Vol. (mL)** | **Initial Weight (g)** | **Final Weight (g)** | **Percent Variance** |
| **1** | 10 mins | 4.0 | 4.0 | 7.0662 | 7.0135 | 0.75% |
| **2** | 10 mins | 4.0 | 4.0 | 7.1540 | 7.1544 | 0.01% |
| **3** | 10 mins | 4.0 | 4.0 | 7.0616 | 7.0596 | 0.03% |
| **4** | 10 mins | 4.0 | 4.0 | 7.1389 | 7.1360 | 0.04% |
| **5** | 10 mins | 4.0 | 4.0 | 7.1588 | 7.1190 | 0.56% |
| **6** | 10 mins | 4.0 | 4.0 | 7.1186 | 7.1167 | 0.03% |
| **7** | 37 Days | 4.0 | 4.0 | 7.1005 | 7.1017 | 0.02% |
| **8** | 37 Days | 4.0 | 4.0 | 7.0339 | 7.1353 | 1.44% |
| **9** | 37 Days | 4.0 | 4.0 | 7.1011 | 7.0559 | 0.64% |
| **10** | 37 Days | 4.0 | 4.0 | 7.0775 | 7.0792 | 0.02% |
| **11** | 37 Days | 4.0 | 4.0 | 7.1679 | 7.1695 | 0.02% |
| **12** | 37 Days | 4.0 | 4.0 | 7.069 | 6.9808 | 1.25% |
| **13** | 105 Days | 4.0 | 4.0 | 7.0714 | 7.0714 | 0.00% |
| **14** | 105 Days | 4.0 | 4.0 | 7.1768 | 7.1774 | 0.01% |
| **15** | 105 Days | 4.0 | 4.0 | 7.0683 | 7.0685 | 0.00% |
| **16** | 105 Days | 4.0 | 4.0 | 7.0977 | 7.0831 | 0.21% |
| **17** | 105 Days | 4.0 | 4.0 | 7.1485 | 7.1479 | 0.01% |
| **18** | 105 Days | 4.0 | 4.0 | 7.1717 | 7.1289 | 0.60% |

### DEMONSTRATION OF CAPABILITY

#### RHODIUM CRYOTUBE FREEZE-THAW TEST PROTOCOL

#### SOP-FRETHAW-003

#### Rev 0

| DATE | DEMONSTRATOR | SIGNATURE |
|---|---|---|

**RHODIUM SCIENTIFIC, LLC**
1300 BAY AREA BLVD, STE B275-5
HOUSTON, TX 77058

MAIN: 210.641.2880

### STANDARD OPERATING PROCEDURE

### EXPERIMENT VERIFICATION TEST

**SOP-EVT-001**

**Rev 0**

PREPARED BY: Dr. R.P. Oates, Program Manager

REVIWED BY: Dr. Heath J. Mills, Chief Scientific Officer

DATE: 12-13-19

This document does not contain technical data as defined in the ITAR, 22 CFR 120.10; or technology as defined under the EAR (15 CFR 730-774).

#### REVISION HISTORY

| REV | CHANGE | SECTION AFFECTED |
|---|---|---|
| 0 | Initial Release, 12-13-2019 | All |

**RHODIUM SCIENTIFIC, LLC**
1300 BAY AREA BLVD, STE B275-5
HOUSTON, TX 77058

MAIN: 210.641.2880

### EXPERIMENT VERIFICATION TEST

**Performed by:**
**Test POC** – Dr. Srivatsan Raman, Assistant Professor, University of Wisconsin-Madison
Phillip Huss, Graduate Research Assistant, University of Wisconsin-Madison

**Directed by:**
**Test POC** – Dr. Heath J. Mills, CSO, Rhodium Scientific, LLC
Dr. R.P. Oates, Program Manager, Rhodium Scientific, LLC

**Test Purpose:**
Determine phage viability and quantity of genomic DNA of *E. coli* post-incubation and freeze in test configurations that match post-flight analytics.

**Test Configuration:**
Rhodium Cryotubes (Part Number: RhCT-0005) were previously filled with 4.0 mL of LB growth media inoculated with *E. coli* and phages and *E. coli* alone during the Phage Evolution pre-flight experiment. This test configuration matches the in-flight configuration.

**Test Protocol:**

1. Remove cryotubes used during biocompatibility tests from the -80°C freezer, one sample set containing *E. coli* and phages combined (ECP samples) in LB media and the second sample set containing *E. coli* only (EC samples) in LB media.

2. Complete in-house standard plaque assay on ECP samples to determine phage viability for incubation times of 1, 2, and 4 hours (timepoint samples in triplicate). Compare results to anticipated minimum for acceptable number of viable phage particles, at least 1 x $10^4$ PFU/mL for 1 hour samples and logarithmic increase at later timepoints.

3. Extract genomic DNA from EC samples for incubation times of 1, 2, and 4 hours to determine quantity of nucleic acids for each sample (timepoint samples in triplicate). Calculate and record DNA concentration and yield. Visually verify DNA quality on a electrophoresis gel. Compare results to minimum concentration of DNA required for downstream analysis (100 ng/μl).

   **Volume of culture:** 1.5 ml for genomic DNA extraction and 1 ml for phage extraction.

**RHODIUM SCIENTIFIC, LLC**
1300 BAY AREA BLVD, STE B275-5
HOUSTON, TX 77058

MAIN: 210.641.2880

**Table 1: ECP Samples – Plaque Assay Results**

| Column A | Column B | Column C | Column D | Column E | Column F |
|---|---|---|---|---|---|
| **Tube Number** | **Incubation Time (at 37$^o$C)** | **# Plaques Observed (µl plated)** | **Dilution** | **Viable Phage Particles (pfu/ml)** | **Cryotube Weight (mg)** |
| **1 $ECP_{1\ hr}$** | 1 hr | 106 (5 µl) | $10^0$ | 2.12E+4 | 7090.9 |
| **2 $ECP_{1\ hr}$** | 1 hr | 105 (5 µl) | $10^0$ | 2.10E+4 | 7065.7 |
| **3 $ECP_{1\ hr}$** | 1 hr | 122 (5 µl) | $10^0$ | 2.44E+4 | 7067.4 |
| **1 $ECP_{2\ hr}$** | 2 hr | 35 (10 µl) | $10^{-5}$ | 3.50E+8 | 7113.3 |
| **2 $ECP_{2\ hr}$** | 2 hr | 18 (10 µl) | $10^{-4}$ | 1.80E+7 | 7106.0 |
| **3 $ECP_{2\ hr}$** | 2 hr | 104 (10 µl) | $10^{-3}$ | 1.04E+7 | 7118.0 |
| **1 $ECP_{4\ hr}$** | 4 hr | 81 (5 µl) | $10^{-5}$ | 1.62E+9 | 7122.3 |
| **2 $ECP_{4\ hr}$** | 4 hr | 120 (5 µl) | $10^{-5}$ | 2.40E+9 | 7055.2 |
| **3 $ECP_{4\ hr}$** | 4 hr | 87 (5 µl) | $10^{-5}$ | 1.74E+9 | 7043.1 |

Notes: 1 ml culture

**Table 2: EC Samples – Genomic Extraction Results**

| Column A | Column B | Column C | Column D |
|---|---|---|---|
| **Tube Number** | **Incubation Time (at 37°C)** | **DNA Concentration (ng/µl)** | **Cryotube Weight (mg)** |
| **1 EC$_{1\ hr}$** | 1 hr | 260.4 | 7086.5 |
| **2 EC$_{1\ hr}$** | 1 hr | 331.1 | 7037.9 |
| **3 EC$_{1\ hr}$** | 1 hr | 264.5 | 7059.3 |
| **1 EC$_{2\ hr}$** | 2 hr | 350.4 | 7064.4 |
| **2 EC$_{2\ hr}$** | 2 hr | 320.5 | 7054.0 |
| **3 EC$_{2\ hr}$** | 2 hr | 407.2 | 7068.6 |
| **1 EC$_{4\ hr}$** | 4 hr | 262.3 | 7064.6 |
| **2 EC$_{4\ hr}$** | 4 hr | 312.5 | 7057.1 |
| **3 EC$_{4\ hr}$** | 4 hr | 318.4 | 7063.8 |

Notes: 1.5 ml culture
Total volume per sample = 80 µl

#### Image 1: EC Samples – Genomic Extraction Gel

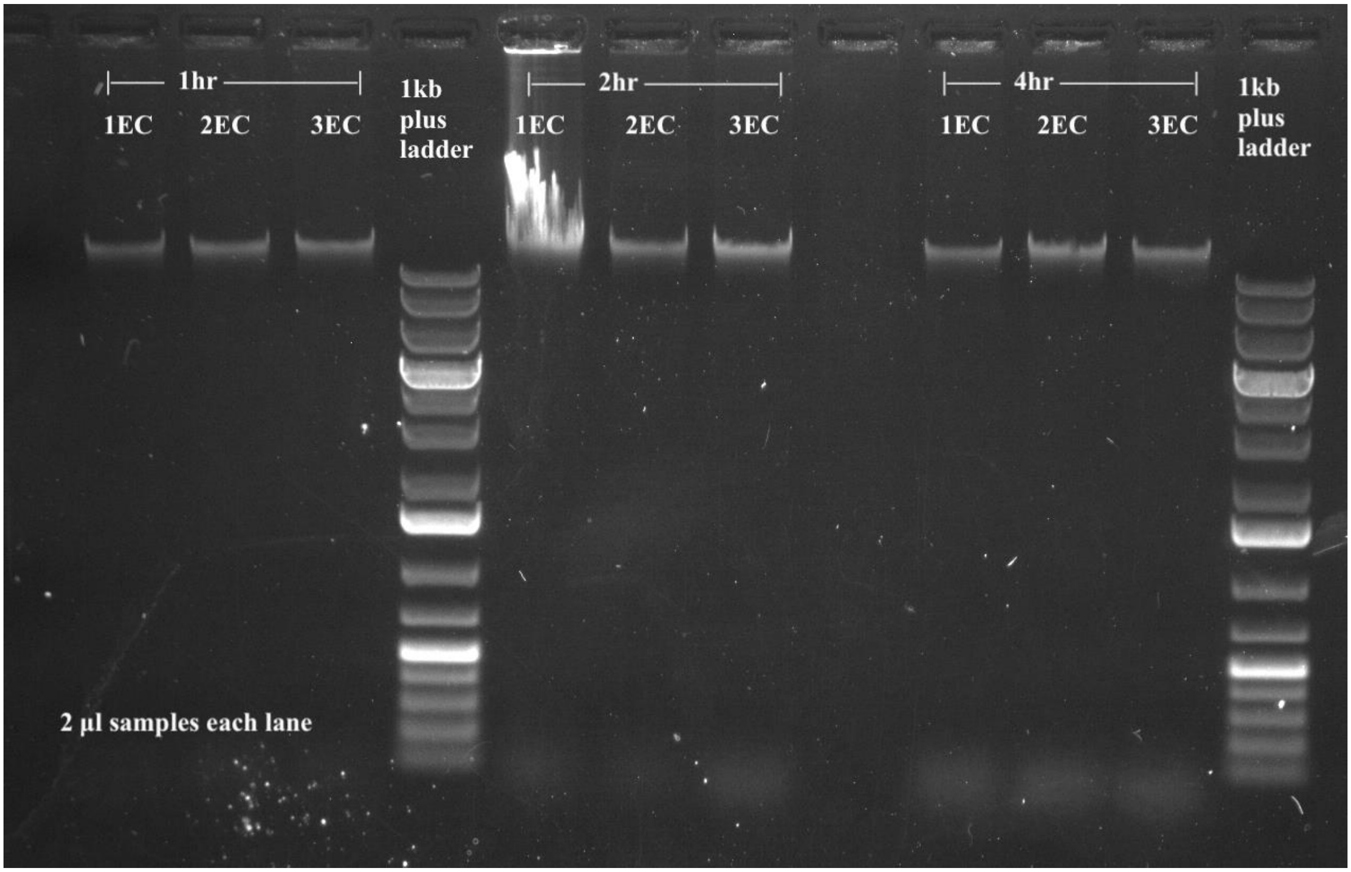

Notes: 120V, 40min, 1%agarose
